# Inter- and intra-individual trajectories of brain sex in different neuroendocrine profiles

**DOI:** 10.64898/2025.12.09.693141

**Authors:** Gloria Matte Bon, Ann-Christin S. Kimmig, Erika Comasco, Birgit Derntl, Tobias Kaufmann

## Abstract

Sex hormone fluctuations modulate structural and functional brain dynamics, yet little is known on how sex steroid levels map onto the expression of sex differences in the brain. Here, we trained machine learning models for brain sex classification based on magnetic resonance imaging regional volumes in cross-sectional data of N = 1090 individuals (50% females, age matched). Applied to densely sampled data of one male and two females in different hormonal states (naturally cycling, oral contraceptive user, pregnancy), we associated inter- and intra-individual fluctuations in brain sex with neuroendocrine profiles. We found lower variation in brain sex across scan sessions in the male compared to the female subjects. Oral contraceptive use was associated with a more female-like brain, while opposite brain sex trajectories emerged across menstrual cycle phases and pregnancy trimesters. Overall, our findings suggest that changes in brain sex capture hormone-related plasticity over time in densely sampled individuals.

## Introduction

Sex differences in brain structure have been traditionally investigated as a binary factor. Binary sex differences have been reported for several regions both in adults and adolescents (*1*, *2*). Such studies often rely on mean comparisons between sexes, overlooking the between subject variability in the population (*3–5*). Interestingly, while findings based on mean comparison are often inconsistent depending on several confounding factors, sex differences in the population variance toward higher male than female variability has been consistently reported for several cortical and subcortical brain regions (*5–7*), pointing toward the need for data-driven approaches that account for population distribution in the study of sex differences (*3*, *4*). It has been suggested that sex differences in the brain may run on a continuum from male-to female-like brain (*8*), in which each individual can manifest on a different position along this continuum, depending on the structure under consideration. The brain is an endocrine organ characterized by the expression of sex hormones receptors. Fluctuations in steroid hormones have been shown to influence brain structure and function (*9–11*), with regions involved in a variety of functions such as emotional regulation, learning and memory showing high expression of sex steroids receptors (*12*). The influence of sex hormones on the human brain starts already in the prenatal period (*13*) and extends throughout the lifespan. This involves a series of genetic and epigenetic mechanisms (*14*), modulating a variety of processes that span from brain plasticity to neurotransmission (*12*, *14*, *15*).

Across the lifespan, different neuroendocrine states have been shown to affect brain structures and function (*11*). Steroid hormones fluctuations during the menstrual cycle have been associated with brain plasticity in multiple cortico-limbic regions (*10*), as well as with changes in cortical network dynamics (*16*). Similarly, oral contraceptives (OC), by suppressing the endogenous sex steroid levels (*17*), have been shown to affect empathy response (*18*) and volumetric changes in different brain structures (*17*). Pregnancy and postpartum on the other hand entail changes in cortical brain anatomy that has been linked to sex hormone levels (*19*, *20*). Such effects, however, have been studied mostly at a univariate level focusing on single structures and functional networks, potentially overlooking the interplay between different regions and networks. Moreover, whether sex hormone fluctuations could affect the expression of sex differences on the female-male continuum in different brain regions is still to be determined.

Machine learning models that classify individuals according to their biological sex have been used to investigate sex differences in the brain in a multivariate setting. Such models have been proposed to offer an insight into the brain sex continuum (*21*, *22*), by returning not only a classification label (male vs female), but also a probability of the position for each individual along a continuum that runs from male-like (coded here as 0) to female-like (coded as 1) brain. Recently, we implemented regional models for brain sex, that classify brain sex based solely on a subset of brain volumes (*23*). Particularly we trained a limbic model, based on regions involved in emotional processing and regulation, and a non-limbic model. We showed sex- and model-specific associations of the obtained probabilities with age, pubertal development (in particular with the biological process of menarche), and female mental health (*24*), highlighting the potential of such models as markers for sex-specific factors.

Here, we aimed to investigate the associations between different neuroendocrine profiles and brain sex. We investigated to what degree brain sex estimates change within the same individual over time and how hormonal states influence such trajectories across adulthood. Building on our prior work, we trained a brain sex classification model based on cortical and subcortical structural MRI volumes of the whole brain, and two regionally constrained models, based solely on limbic or non-limbic volumes. Training was based on cross sectional data derived from the Human Connectome Project Young Adult (HCP-YA) (*25*), the Human Connectome Project Aging (HCP-A) (*26*) and the Queensland Twin IMaging (QTIM) (*27*) cohorts. We applied these models to densely sampled brain imaging data from one 23 years old female tested 30 consecutive days while naturally cycling to cover an entire menstrual cycle (28andMe), the same female recorded one year later for 30 consecutive days while taking hormonal contraceptive (28andOC) (*16*, *28*), one 38 years old female tested 26 times to cover an entire pregnancy, starting with pre-conception until 2 years postpartum (Maternal Brain Project) (*29*, *30*), and one 26 years old male recorded 40 times over 30 consecutive days every 12-24 hours (28andHe) (*31*, *32*). These yielded estimates of brain sex for each individual and time point across the dense sampling. We then investigated the trajectories of brain sex estimates across time points and correlated them with hormonal levels. Our hypothesis was that brain sex predictions vary across time and between individuals in different hormonal states. Specifically, given hormonal changes across pregnancy and across the cycle, we expected greater variability across time in females compared to the male participant, and that associations between brain sex and hormones will be stronger for the limbic compared to the non-limbic model, due to the high concentration in hormonal receptors in limbic structures (*11*).

## Results

Our training of global and regional models in cross-sectional data yielded a limbic, non-limbic and whole brain model of brain sex. Specifically, to reduce variance due to difference in brain size between females and males that could influence our classification models, we combined in our training sample data from the HCP-YA, HCP-A and QTIM samples and we matched each female participant to one male based on i) a difference in estimated total intracranial volume (eTIV) less than 3%, ii) a difference in age of maximum 1 year and iii) a difference in Euler number (a proxy of image quality) (*33*) of less than 1 standard deviation from the mean. The final training sample was composed of N = 1090 participants (50% females, age range: 19-100 years old, age mean ± SD = 36.4 ± 18.6), with no significant differences in age (t = -0.016, p = .987), eTIV (t = -0.702, p = .483) and Euler number (t = -0.114, p = .909) between sexes. Supplementary Figure S1 shows the final distribution of eTIV for the selected sample compared to the general population. The selected sample included values close to the maximum and minimum of the distribution of the full sample and extended over the mean and median values of the general population for each sex, limiting potential effects due to biased selection of individuals with smaller or larger brain only, while controlling for eTIV by removing the significant sex difference for this confounding effect in the general population (t = -39.52, p < 2x10-16).

All models achieved good cross-validation performances in our matched sample, with an area under the curve (AUC) greater than 0.80 for all models (limbic = 0.817, non-limbic = 0.847 and whole brain = 0.880) (Figure 1a). In accordance to our previous works on adults and adolescents (*23*, *24*), model performance was similar between all models despite the different number of features, with the limbic model having nearly 1/3 of the features compared to the whole brain model (limbic: 160 features, non-limbic: 333, whole brain: 493), pointing toward the relevance of limbic volumes in capturing sex differences (see Supplementary Table T1 for the detailed list of limbic features). Furthermore, feature contributions were correlated between the regional models and the whole brain model, suggesting that the models are robust (Supplementary Figure S2).

**Figure 1.**
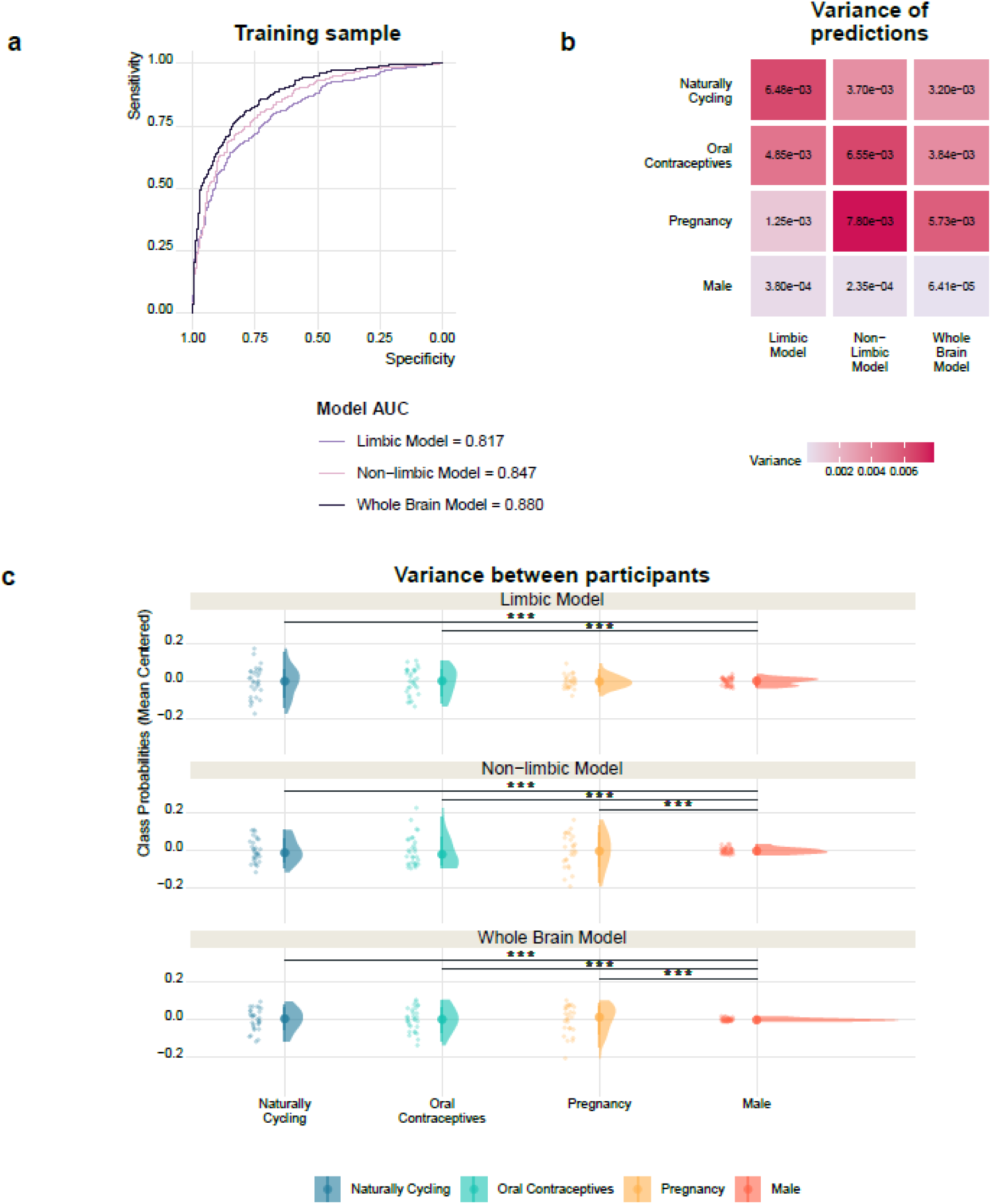
Brain sex predictions showed inter- and intra-individual variability. **a)** Cross-validation in cross-sectional data yielded high AUC for all models. **b)** Variance of class probabilities for each model and subject. Darker color corresponds to higher variance of class probabilities. **c)** The male participant showed significant lower variance in brain sex predictions compared to female participants for all model except the limbic model during pregnancy. Class probabilities are centered around the mean for each group. *** p < .003 (Bonferroni-corrected threshold)

When compared to models trained on a sex-balanced sample matched only by age and Euler number (N = 2310, 50% females, age mean ± SD = 34.9 ± 17.5) with different eTIV correction methods, our eTIV-matched models achieved comparable AUC to models in which eTIV was accounted for through eTIV regression (limbic = 0.807, non-limbic = 0.871, whole brain = 0.896) and power-corrected proportion methods (PCP) (*34*) (limbic = 0.769, non-limbic = 0.793, whole brain = 0.833) (Supplementary Figure S3, panel b-c), while lower performances compared to the models trained on raw features (limbic = 0.927, non-limbic = 0.917, whole brain = 0.939) were found for all eTIV-corrected models (Supplementary Figure S3a) independently from the sample size. These findings point toward eTIV correction rather than the reduction in sample size as responsible for the lower performances compared to models trained on raw features. Furthermore, class probabilities in the eTIV-matched sample were significantly correlated for all models independently from the eTIV correction method (raw features, eTIV regression and PCP, Supplementary Figure S3d-f), with a Cohen’s Kappa agreement measure spanning between moderate to substantial (Supplementary Table T2). On the other hand, models trained on the participants excluded from the matching procedure and therefore highly biased toward the lower and upper end of eTIV distribution in the overall population yielded AUC of over 0.98 for all models (limbic = 0.985, non-limbic = 0.990, whole brain = 0.992) (Supplementary Figure S4), achieving almost perfect classification and suggesting highly biased predictions. Taken together, these results confirm our eTIV-matching approach as a robust method to control for head size in the training sample. When externally validating the models in densely sampled data, all models were largely able to classify sex in each sample correctly for the majority of timepoints (Supplementary Figure S5b), as shown by class probabilities lower than 0.5 (binary labeled as male) for all timepoints of the male individual, and greater of 0.5 (binary labeled as female) for most of the timepoints for the female participants (Supplementary Figure S5b). Yet we observed variance in class probabilities across time and between subjects, reflecting daily variation in brain structures captured by the models. Levene’s test allowed us to test for equality of variances on the centered distributions. Interestingly, the male subject had consistently significant lower variances for all models (limbic: F = 17.216, p = 2.133 x 10^-9^; non-limbic: F = 15.296, p = 1.612 x 10^-8^; whole brain: F = 18.753, p = 4.421 x 10^-10^) compared to the female subjects in all hormonal states, except for the limbic model in the pregnant woman that had similar variance. Figure 1b and c depict the variance of class probabilities and the respective class probabilities, centered to the mean for direct comparison between samples. Supplementary Figure S5 depicts the distribution of class probabilities in the different samples. Supplementary Figure S6 shows class probabilities for the male individual divided by morning and evening sessions. No significant difference between morning and evening sessions was found for any of the models for the male participant (limbic: t = -0.339, p = .739; non-limbic: t = -1.481, p = .155; whole brain: t = -1.481, p = .155). Supplementary analysis in an independent sample composed of 30 healthy adults (50% females) that underwent 10 scans across one month (one scan every three days approximately) (*35*) revealed similar patterns of higher variance of brain sex estimates in females compared to males, with a significant effect for the limbic model (p = .025) (Supplementary Figure S7a). To further confirm this patterns, we applied our models also to data from the Neurocycle+ study (*36*, *37*), a dense sampling study in which one naturally cycling female, one female taking oral contraceptives and one male individual were tested 25 times over one month. We found a similar, general pattern of higher variance in the female participants, particularly the naturally cycling, compared to the male for the limbic and whole brain model, while a significant difference between oral contraceptive user and naturally cycling was found in the non-limbic model (Supplementary Figure S7b). However, the non-limbic model classified the naturally cycling female as male-like brain for all timepoints. Although class probabilities were still significantly higher than for the male subject (t = 3.127, p = 0.003), the probabilities were below 0.5 (Supplementary Figure S8). Therefore, the significant difference in variance between oral contraceptive user and the naturally cycling female for the non-limbic model might stem from the low class probabilities in the naturally cycling female toward male-like brain.

To further explore whether patterns of higher variance in females were present already in the cross-sectional training sample, we performed a Levene’s test between sexes on the class probabilities obtained with the cross-validation in the training sample. After correcting for the three models, no significant difference was found in the variance of cross-sectional class probabilities between males and females (limbic: F = 4.681, p = .031; non-limbic: F = 0.490, p = .484; whole: F = 0.898, p = .344), further suggesting the involvement of the temporal component in determining this effect in the densely sampled individuals.

Investigating the source of these multivariate brain sex distributions, we turned to the univariate level and assessed how much single brain regional volumes varied over time in the naturally cycling female compared to the male subject, especially given earlier reports of higher structural variance in boys than girls (*5*). Interestingly, coefficients of variation for single brain regions appeared overall more similar between subjects than what was observed at the multivariate level (Supplementary Figure S9), suggesting that lower variance in class probabilities for male brain sex over time was not simply attributable to lack of temporal variation in anatomical data.

Given the impact of oral contraception (OC) on neuroendocrine balance, we tested how OC use affected brain sex, by comparing intraindividual changes in brain sex when naturally cycling and when taking OC in the 28andMe. Our findings showed increased certainty in predictions when taking oral contraceptives, toward higher class probabilities for all models (more female-like brain) (Figure 2a). Such increase was significant only for limbic estimates (t = -3.129, p = .004), while no significant difference was found for the non-limbic (t = -1.399, p = .172) or whole brain (t = -1.194, p = .242) models. Interestingly, a similar inter-individual pattern of higher class probabilities in the OC user was found in the NeuroCycle+ data, where the participant using OCs showed significantly higher class probabilities for the non-limbic and whole brain models compared to the naturally cycling individual (Supplementary Figure S10). However, the male-like predictions for the naturally cycling participant drive this difference in the non-limbic model.

**Figure 2.**
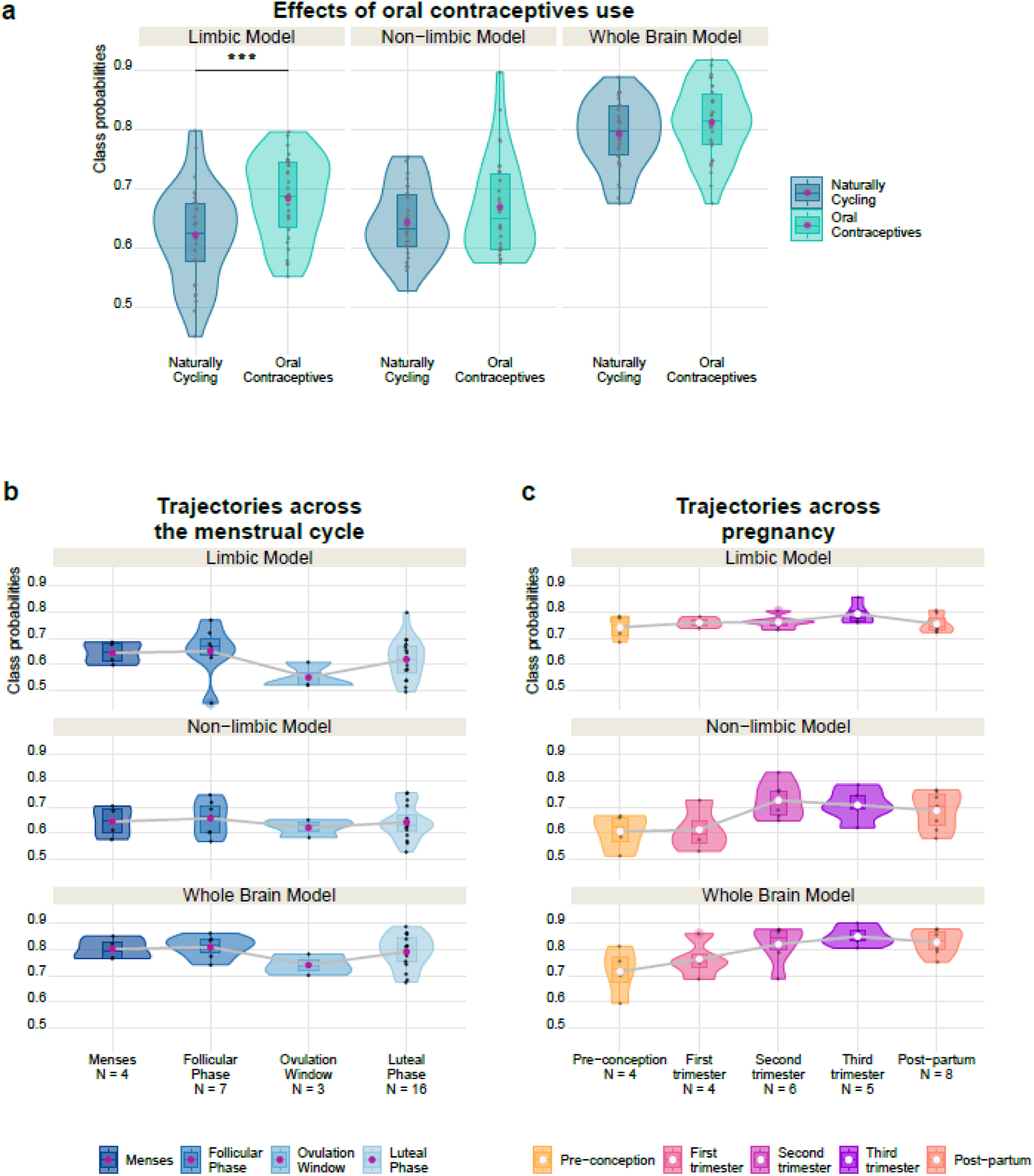
Intra-individual variation in class probabilities across hormonal states. **a)** Oral contraceptive use is associated with significantly increased class probabilities toward more-female like brain for the limbic model. No significant effect was found for the non-limbic or whole brain model. **b)** Class probabilities showed a minimum toward more male-like brain around ovulation for all models. N indicates the number of days assigned to each phase based on the information available from the data provider **c)** Across pregnancy, class probabilities showed trajectories toward an increase in class probabilities and more female-like brain that partially reverted post-partum. N indicates the number of data points available for each trimester. *** p < .017 (Bonferroni-corrected threshold).

We then visually investigated the individual trajectories across hormonal states, focusing on the trajectories across the menstrual cycle’s phases and across pregnancy trimesters. Our results showed consistent trajectories of class probabilities across menstrual cycle phases between models, with a negative peak toward more male-like brain around ovulation for all models (Figure 2b). Interestingly, the limbic model showed the strongest dip during ovulation, while the non-limbic model showed less variability across phases. Of notice, each phase has a different length in days, resulting in different numbers of observations. Moreover, since the first session was blinded for cycle phase, the experimental sessions spanned across two consecutive menstrual cycle, covering the period between the luteal phase of cycle 1 and the mid-luteal phase of cycle 2. Therefore, the luteal phase includes data from two different cycles. Trajectories across the two cycles are shown in Supplementary Figure S11.

To further investigate the dynamics across the menstrual cycle, we look at the trajectories in the naturally cycling female of the NeuroCycle+ sample and we found consistent trajectories across menstrual cycle phases between models, with an increase in class probabilities toward more female-like brain in the second part of the cycle (ovulation, luteal phase) across two consecutive cycles, confirming intra-individual fluctuations across cycle phases (Supplementary Figure S12a). It is important to highlight that the NeuroCycle+ study included less timepoints over 30 consecutive days compared to the 28andMe, resulting in different numbers of observations for each phase and some missing days in the cycle.

To explore such dynamics in bigger sample sizes, we ran a supplementary analysis in three longitudinal samples: the HNU, the Hormonal Health Study (HHS) (*38–40*) and the sample described in Wright et al. (*41*, *42*). The HHS and the Wright et al. sample collect longitudinal data of female participants at 3 timepoints, each capturing a different cycle phase (menses/early follicular (EFP), ovulation window/late follicular (LFP) and mid-luteal phase (MLP)). For the HNU sample, we assign sessions for each female participants to EFP, LFP and MLP based on the days from the beginning of the last menstrual period. Day intervals were established based on the Wright et al. sample for consistency as early follicular (days 1 to 4), late follicular (days 10 to 12) and mid-luteal (days 20 to 22). We found a consistent inter- and intra-individual variation across menstrual cycle phases in all samples (Supplementary Figure S12b-d), further highlighting the expression of brain sex as dynamic within each individual across neuroendocrine states. It is important to highlight that while similar patterns between models emerged for the HNU sample (Supplementary Figure S12b), in which multiple scans could fall in the same cycle phase for each individual, no general differences between cycle phases emerged across the HHS and the sample from Wright et al. (Supplementary Figure S12c-d), in which each participant was tested only once for each phase, suggesting that more densely collected data are necessary to depict the full picture of brain dynamics across the menstrual cycle.

When plotting class probabilities trajectories across pregnancy trimesters, starting with pre-conception until postpartum, a pattern of increased class probabilities across pregnancy was found for each model (Figure 2c). Class probabilities increased toward more female-like with pregnancy progression, with a maximum around the third trimester for the limbic and whole brain models and in the second trimester for the non-limbic model, that partially reverted in the postpartum period for all models. While a constant increase across trimesters was found for the whole brain, the non-limbic class probabilities showed a steep increase between the first and the second trimester. Interestingly, the limbic model showed less variability in this case. Nevertheless, the single subject design does not allow for statistical testing of differences between phases neither for the naturally cycling nor for the pregnant individual.

Finally, we investigated the correlation between class probabilities and hormonal concentration. Given the high number of tests, none of them survived Bonferroni correction for multiple comparisons yet we report the few nominally significant correlations between brain sex and hormonal levels in the naturally cycling, pregnant and male participants. Estradiol levels were negatively correlated with class probabilities for the limbic model (r = -0.413, p = .023) and whole brain model (r = -0.381, p = .038) in the naturally cycling woman (although both failed to pass correction for multiple comparisons), while no nominally significant correlation was found for the non-limbic model or other hormones such as progesterone, LH and FSH (Figure 3a, e, f). For pregnancy, we found a nominally positive correlation with progesterone for the whole brain model (r = 0.597, p = .040) when associating data of the first, second and third trimester (Figure 3c, g). Interestingly, a nominally positive significant association with estradiol levels toward more female-like brain was found in the male participant (r = 0.427, p = .019) (figure 3d, h). When dividing the data by time of the day, such association was found only for the evening sessions (r = 0.640, p = .010) (Supplementary Figure S13). Finally, no significant associations were found for the female participant when taking oral contraceptives.

**Figure 3.**
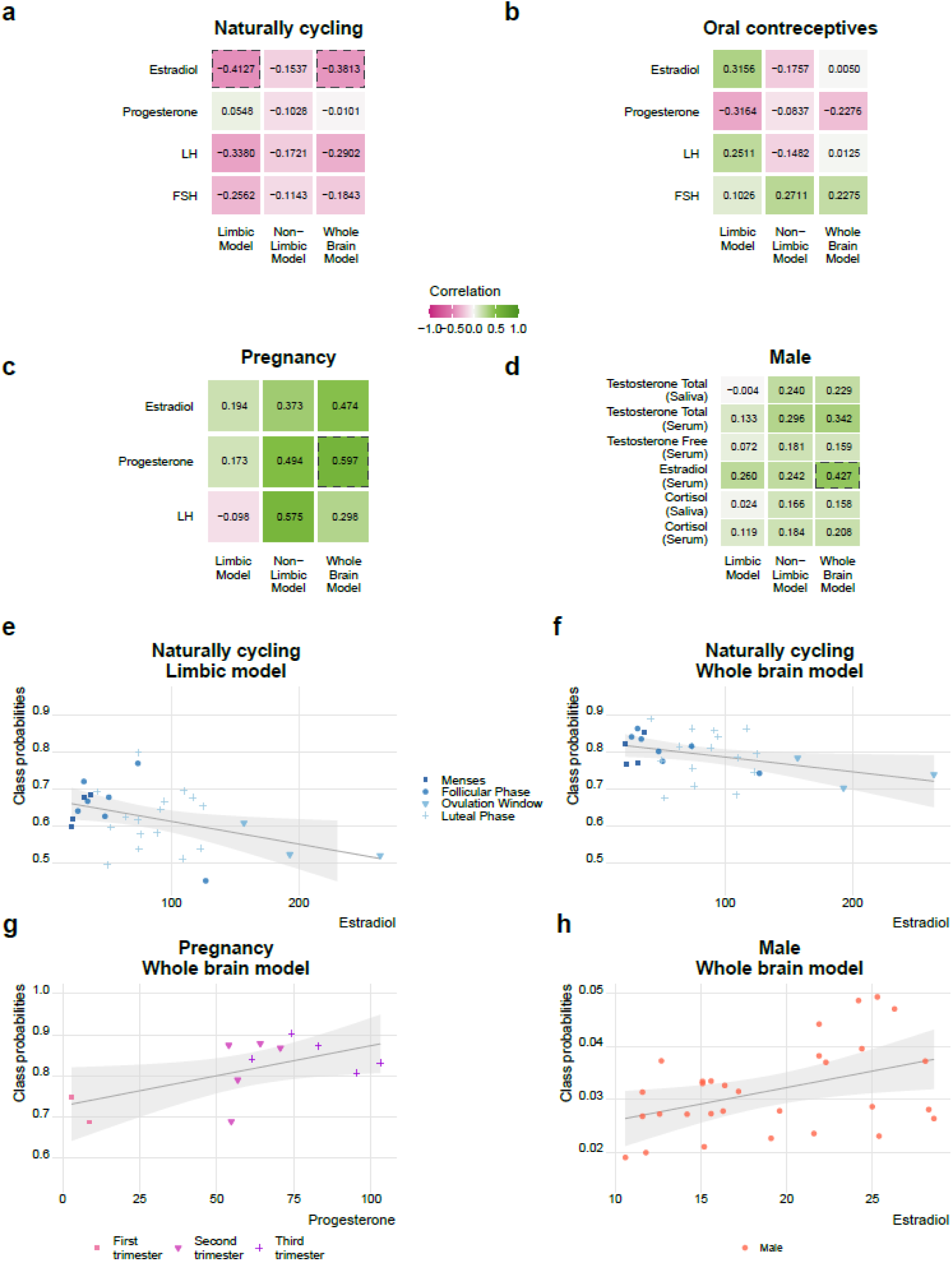
Correlation of hormonal levels with class probabilities per participant. Class probabilities are negatively associated with estradiol levels in the naturally cycling participant for the limbic and whole brain model **(panel a, e and f)**, while no significant association was found in the same participant when taking oral contraceptives **(b)**. Across pregnancy trimesters, a positive association between whole brain estimates and progesterone levels was found **(panel c, g)**. Finally, a positive association between estradiol levels and whole brain estimates was found in the male subject **(d, h**). However, none of the correlations survived Bonferroni correction for multiple comparisons. Pink color in the heatmap indicates a negative correlation (decrease in class probabilities) while green indicates a positive correlation (increase in class probabilities). Dashed lines: uncorrected p < .05.

Based on the nominally significant correlation between limbic and whole brain estimates with estradiol levels, we explored the temporal dynamics between class probabilities and estradiol using vector autoregressive (VAR) models. Based on Pritschet et al. (*16*), we considered second-order VAR models for consistency with previously reported effects in the same data, and we further explore this association with third-order VAR based on the model information criteria. We found significant associations of limbic brain sex with estradiol in both models, particularly with the previous day (lag 1, t = -2.421, p = .0238) for the second-order and the previous day (lag 1, t = -3.477, p = .00238), two days before (lag 2, t = 3.082, p = .00588) and three days prior (lag 3, t = -2.307, p = .03190) for the third-order VAR (Supplementary Figure S14). Significant Granger causality test for third-order VAR (F = 4.325, p = .00987) confirmed the temporal association between estradiol and limbic estimates.

## Discussion

This work illustrates inter- and intra-individual changes in brain sex estimates based on different sets of structures across different neuroendocrine profiles. Our findings showed greater general variability of brain sex predictions in densely sampled data of female participants compared to the male participant across time (i.e., scan sessions) for all hormonal states, and different class probability dynamics in different neuroendocrine profiles.

Class probabilities across timepoints showed lower variance in the male participant compared to both females, independently from the hormonal state. These results were confirmed in supplementary analyses in two independent samples. Of notice, when considering single volumes, coefficients of variation were comparable between the naturally cycling female and the male participant. Greater male than female variability in single brain structures has been reported in the literature across different age periods (*5*, *6*, *43*). However, such studies are mostly focused on cross-sectional univariate differences across individuals in different age ranges. Here, we investigated the variance associated with longitudinal multivariate estimates in a short period of time in the same individual, showing that sex class probabilities based on brain structural MRI change in the same individual across timepoints. Interestingly, from a functional perspective, previous studies comparing connectivity dynamics in the same densely sampled individuals found greater variability in the male subject compared to the naturally cycling female, suggesting different patterns of variability for brain structure and function even in short periods of time (*32*, *44*). It must be highlighted that the timeframe of the data highly differs between the pregnant participant and the others, pregnancy related data being collected over almost two years, compared to the 30 days of the menstrual cycle, oral contraceptive use and male participant data. However, class probabilities showed comparable variance across time between pregnancy and menstrual cycle data, suggesting the involvement of other factors than time in determining this variance, potentially pointing toward the neuroendocrine modulation as the driving source.

When investigating the effect of different hormonal profiles in the same individual, we found increased class probabilities toward a more female-like brain when using oral contraceptives for all models, with a significant effect for the limbic model. These findings were partially validated in an independent dense sample, where inter-individual differences in brain sex were found, with higher class probabilities reported in the participant taking oral contraceptive compared to the naturally cycling participant. Limbic regions are highly dense in sex steroid receptors that can mediate structural changes in the brain (*45*, *46*). Studies investigating the effects of oral contraceptives have been so far focused on single structures and cross-sectional data, with only few studies analyzing longitudinal data with different time intervals (*17*). Oral contraceptives have been reported to have different effects on the brain, having been associated with both an larger and smaller gray matter volume depending on the structure under consideration and the type of oral contraceptive (*17*). The increase in class probabilities may be related to the high concentration of synthetic sex hormones associated with oral contraceptive use, and in particular the high concentration of the progestin levonorgestrel (*47*). Here, by investigating intra- and inter-individual effects on a multivariate measure of brain sex, we explore the effects of such hormones on sex differences when considering the whole sex mosaic.

Fluctuations in several brain volumes across the menstrual cycle and across pregnancy have been previously reported in the literature (*10*, *16*, *19*, *36*, *46*, *48*). However, studies investigating how changes across the menstrual cycle influence the expression of sex differences in the brain are scarce and mostly in a cross-sectional setting, focusing on group difference between NC and OC use vs males (*49*). Likewise, sex differences in brain structure and function in the context of pregnancy and parenthood are still largely under investigated, with only few studies investigating such dynamics in males (*50*, *51*). By looking at the trajectories of brain sex across these periods, we aimed to investigate the effects of hormonal states on the predictions in a longitudinal setting. We found consistent trajectories between models across both the menstrual cycle and the pregnancy. These findings were further confirmed by supplementary analysis in independent samples, showing inter- and intra-individual differences in brain sex across menstrual cycle phases, supporting the notion of brain sex as a dynamic factor influenced by neuroendocrine states.

The negative peak toward more male-like brain around ovulation during the menstrual cycle was in line with our findings of negative correlation of class probabilities with estradiol levels, that peaks around ovulation (nominally significant but not passing corrections), suggesting that the models may to some extent capture plasticity related to sex hormones exposure. Associations between whole brain structural dynamics and gonadal hormones fluctuations have been reported in densely sampled female participants with different hormonal profiles (*36*), suggesting an effect of hormonal fluctuations on brain structure even in a short period of time. However, we cannot exclude the effects of other factors and hormones not analyzed here, such as testosterone, which peak overlaps with the estradiol one. Likewise, because no data on hormonal exposure was available in the cross-sectional training set, we were not able to take this into account for brain sex modelling. However, in including a wide age range across the lifespan we have diminished the impact of selection bias for a specific hormonal phase. Moreover, individual fluctuations between menstrual cycles in the same individual can also affect the predictions. Here, the experimental sessions partially covered two consecutive cycles, potentially affecting the predictions. When dividing the luteal phases based on the cycles (cycle 1 and cycle 2), the overall dynamics rest largely unchanged, with the negative peak around ovulation (Supplementary Figure S11). However, differences in class probabilities between the two luteal phases emerged, suggesting intra-individual variation in brain sex estimates, possibly influenced by intra-individual fluctuation in hormonal levels between cycles. Nevertheless, in light of the nominally significant negative correlation with estradiol levels, intra-individual fluctuations between cycles are unlikely to entirely explain the effects toward more male-like brain around ovulation. Temporal dynamic association between limbic estimates and estradiol levels investigated through VAR models further support this hypothesis. Interestingly, the oscillation between negative and positive sign between lags was found also by Pritschet and colleagues (*16*), suggesting a role of estradiol in regulating not only functional but also structural brain dynamics across the menstrual cycle. Interestingly, estradiol levels showed the opposite trend in the male participant, correlating with an increase in class probabilities toward more female-like brain (although failing in passing correction for multiple comparisons).

Across pregnancy, all models showed an increase in class probabilities toward more female-like brain that partially reverts in the postpartum period. U-shaped trajectories of gray matter volumes for several structures across pregnancy were reported in the literature, with a local minimum in the third trimester that partially reverted post-partum (*30*, *52*). Furthermore, changes in gray matter volume have been previously related to both estradiol (*30*, *53*) and progesterone dynamics across pregnancy (*30*). Here, we found opposite trajectories (increase in class probabilities that partially reverted post-partum), nominally correlated with increasing levels of progesterone. The increase in class probabilities toward more female-like brain could be partially attributed to the gray matter volume loss across pregnancy (*48*). Differences in total intracranial volume between sexes are consistently reported in the literature, with males showing larger brain volumes compared to females, potentially affecting the brain sex predictions (*54*, *55*). Our training sample was matched based on eTIV, ensuring no significant difference in eTIV between sexes. To further control for other processes in our sample that could involve volume loss and result in a biased classifier toward a more female-like brain (e.g., aging), we checked for an interaction effect between sex and age on eTIV in our training sample (Supplementary Figure S15) and found no significant differences (t = 0.128, p = .898). Moreover, a supplementary analysis training the models on a subsample with an age range between 19 and 49 years, closer to the one of the densely sampled participants, yielded similar estimates for all participants as well as similar trajectories both across the menstrual cycle and pregnancy (Supplementary Figure S16). Therefore, the trajectories toward more female-like brain found across pregnancy are unlikely to be attributed uniquely to the volume loss in the pregnant female. Moreover, many of the structures reported to undergo gray matter volume loss during pregnancy are included in the limbic model (*30*). Of notice, the limbic model in pregnancy showed lower variance in the predictions, further supporting the notion that the shown effects are not dependent uniquely on gray matter volume loss.

Changes in brain structure from pre-conception to early post-partum have been previously linked to the mean levels of estrogens during pregnancy, particularly during the third trimester (*53*). Interestingly, our results showed a nominally significant correlation with progesterone levels during pregnancy trimesters in the whole brain model, in line with the univariate findings in the same data linking structural brain dynamics to the significant rise in steroids hormone concentration over pregnancy (*30*). Different factors could be responsible for the discrepancy in the findings on the correlations of different hormones with brain changes reported in the literature, including different collection methods for the hormones considered (urine samples collected at home versus blood draw before the MRI scanning) and sample size differences. Of notice, while previous studies linked estradiol levels during pregnancy to MRI scans carried out pre-conception and post-partum (*53*), the hormonal levels were here collected on the same day of the MRI scanning, allowing a direct measure of the neuroendocrine profile for each study appointment. Nevertheless, the densely sampled nature of the data hinders any generalization on the direct effects of hormones in determining such changes. Furthermore, other factors not accounted for in this study such as the hormonal stimulation carried out to achieve pregnancy via in vitro fertilization (IVF) might influence the results.

Despite the limited sample size, the trajectories across neuroendocrine profiles taken together allow us to speculate on possible underlying mechanisms. The negative correlation of class probabilities with estradiol although only nominally significant, together with the negative peak of predictions around ovulation (corresponding to the estradiol peak) and the temporal association between limbic estimates and estradiol concentration in the naturally cycling female suggest an effect of estradiol toward lower class probabilities and more male-like brain, while the trajectories across pregnancy point toward an effect of the combination of high levels of progesterone and estradiol in shaping the increase in class probabilities toward more female-like brain. This hypothesis is also in line with our findings in the OC user. Combined OC administration induces a relatively steady level of synthetic hormones in different combination, with a smaller dose of estrogens and a higher dose of progestin, that interfere with the normal endogenous production of the endogenous analogue hormones (*47*), resulting in the combination of high doses of progestogens and estrogens. Interestingly, while the endogenous production of progesterone was suppressed by OC administration, endogenous estradiol dynamics were unaltered (*16*). However, the complex interaction between high doses of synthetic and endogenous hormones is likely to exert a stronger effect than the endogenous production of estradiol alone (*47*) and to be held responsible for the findings. Taken together, our results suggest opposite effects of estradiol alone and its combination with progesterone on brain sex predictions in females. Nevertheless, different hormonal sensitivity as well as different baseline positions along the brain sex continuum could result in inter-individual differences in the trajectories. While the reduced sample size in the dense sample and the lack of repeated measures for each phase of the cycle in bigger samples do not allow to infer on the direction of the effects at higher population level, our findings highlight the presence of inter- and intra-individual differences in the trajectories of brain sex across cycle phases. Few methodological limitations need to be addressed. Although the dense sampling study design allows to investigate intra-individual trajectories, it poses limitations regarding the generalizability of findings, particularly regarding inference on sex differences and intra-versus inter-individual variability. The lack of longitudinal data regarding hormonal profiles in big samples constitutes one of the major limitations in the study of neuroendocrine influences on these models. Even when neuroendocrine factors such as menstrual cycle phases are collected, the lack of repeated measurement in the same phase does not allow to account for day-to-day variability, limiting the investigation of dynamic processes. Dense sampling approaches, by providing extensive data on single or few participants, constitute a major resource to investigate intraindividual changes and short-time longitudinal dynamics. However, with the data available, we are unable to generalize our findings at a higher population level. Thus, future research integrating short-time longitudinal assessments with neuroendocrine information, such as circulating hormonal levels, menstrual cycle phase or oral contraceptive use, in large sample studies is needed to further generalize these effects.

Moreover, the few participants and the longitudinal nature of the data did not allow to apply any direct eTIV correction methods in the test samples. However, the matching procedure in the training sample ensured independence of class probabilities from eTIV, with no significant eTIV difference reported between the two sexes. Importantly, eTIV distribution in the densely sampled participants overlapped with the matched training sample (Supplementary Figure S17). It is worth to mention that the naturally cycling participant had a bigger eTIV than the male individual. Nevertheless, both individuals were correctly classified according to their biological sex, suggesting that the matching procedure was effective in limiting the bias due to bigger eTIVs in males compared to females. Different eTIV correction methods have been previously shown to have an effect on brain sex estimates (*23*, *24*, *54*, *56*). Here, the longitudinal nature of the data collected in a very short period, where eTIV fluctuations are very small, limited the option for traditional methods for confound controls (e.g., residuals approach, PCP). By applying a stringent matching procedure based on the matching procedure developed by Wiersch and colleagues (*56*) and previously validated for our models in an adolescents sample (*24*), we were able to limit potential effects due to eTIV in our models. Of notice, while reducing the overall sample size and consequentially potentially reducing the statistical power of the analysis, eTIV matching is the only method to completely remove sex-related eTIV variation in the sample (*34*) without touching the features. In line with the literature (*34*), the comparisons with residuals and PCP correction in the training sample confirmed highly consistent results, further validating the matching procedure approach for eTIV correction in multivariate analysis. Moreover, the eTIV distribution in the matched sample largely overlapped with the distribution in the full sample for each sex reaching almost the maximum and the minimum eTIV values, ensuring that the final training sample was to some degree representative also of individuals with larger or smaller eTIV for both sexes (Supplementary figure S1).

Finally, the lack of harmonization to control for between scanner effects in the pregnancy data might affect the results. However, a validation scan performed within 24 hours from the end of the study in the original work ensured highly consistent measures between scanners, indicating that scanner effects are unlikely to explain the results (*30*).

Overall, our findings highlight the potential effect of neuroendocrine profiles on brain plasticity, influencing not only structural dynamics at a univariate level, but also multivariate estimates of brain sex derived from structural data.

## Methods

### Participant selection

Study data relied on a combination of multiple open data. Each sample obtained the approval by the respective local ethics committee.

The training sample was composed of a combination of participants from Human Connectome Project Young Adults (HCP-YA) (*25*), the Lifespan Human Connectome Project Aging (HCP-A) (*26*) and the Queensland Twin IMaging study (QTIM) (*27*). After the exclusion of 14 participants from the HCP-YA, 20 participants from QTIM and 22 participants from the HCP-A due to data quality issues, we selected participants with an age range between 19 and 100 years, leading to a total of N = 2744 subjects (58% females, HCP-YA = 1098, QTIM = 934, HCP-A = 712). The age range was selected based on data availability, including diverse stages of life with the aim to cover diverse hormonal profiles (e.g. including post-menopausal females, as well as individuals that had previous pregnancy, etc.). To remove the effects due to multiple scanners for image acquisition, we harmonized the data using the *neuroCombat* harmonization for multi-site imaging data (version 1.0.13) (*57*) across six sites (1 site for HCP-YA, 1 for QTIM and 4 for HCP-A), accounting for sex and age as covariates. To guarantee that our final training sample was balanced according to sex and at the same time control for confounders such age and brain size, we implemented a multistep matching procedure (*24*, *56*). After dividing participants according to their biological sex assigned at birth, each female was matched with one male, so that 1) the maximum absolute difference in age was one year, 2) the maximum absolute difference in the harmonized estimated total intracranial volume (eTIV) was less than 3% and 3) the absolute difference in image quality was within 1 standard deviation. If more than one male participant fitted the criteria, the participant with minimum difference in eTIV was selected as best match. The final training sample was composed by 1090 participants (50% females, age range: 19-100 years old, mean ± SD: 36.4 ± 18.6, subsample composition: HCP-YA = 355, HCP-A = 351, QTIM = 384), with no significant difference in age, eTIV or image quality between sexes. To further control for age effects we carried out a supplementary analysis in a subsample of the matched participants with an age range between 19 and 49 years (N = 844, 50% females), closer to the age range of the densely sampled participants.

Since eTIV matching reduces the sample size, potentially affecting the predictions accuracies of our models, in a supplementary analysis we compared the models trained in the eTIV-matched sample with models trained in a sex-balanced sample matched only for age and image quality as described in point 1 and 3 of the matching procedure, resulting in a sample of N = 2310 (50% females, age mean ± SD = 34.9 ± 17.5). To further disentangle in this sample the effects due to the sample size and the eTIV correction, we compared models trained on 1) raw features without any eTIV correction method; 2) residuals after eTIV was regressed from each feature prior to model training; 3) PCP corrected features (*23*, *34*). To further confirm the efficacy of the matching procedure in reducing bias related to eTIV, we carried out a supplementary training of the models on a sample composed by the participants excluded from the matching procedure (N = 1654, 63% females, age mean ± SD = 33.8 ± 16.2), in which eTIV differences between sexes were accentuated.

To investigate individuals’ trajectories across different hormonal states, we used independent data derived from three dense sampling studies as external validation samples: the 28andMe and 28andOC, the 28andHe and the Maternal Brain Project (*16*, *28–32*). The 28andMe (*16*, *28*) consists of data from one naturally cycling (NC) female participant (23 years old, history of regular menstrual cycles occurring every 26-28 days) recorded for 30 consecutive days to cover the length of an entire menstrual cycle. In the 28andOC, the same participant underwent the same protocol again one year later for 30 consecutive days while taking oral contraceptives (OC, 0.02 mg ethinyl-estradiol, 0.1 mg levonorgestrel, Aubra, Afaxys Pharmaceuticals, started 10 months prior to the start of data collection). Study protocol for both studies included behavioral assessments, blood collection for endocrine assessments and MRI acquisition (*16*). Cycle phases were determined based on hormonal levels. The 28andHe (*31*, *32*) includes one 26 years old participant who underwent testing every 12 or 24 hours for 30 consecutive days for a total of 40 sessions. Sessions were scheduled in the morning (7 a.m., day 1-15) and/or in the evening (8 p.m., day 11-30) every day. At each session, the participant underwent behavioral testing, MRI scanning as well as saliva and blood collection for endocrine assessments. In days with morning and evening sessions, blood was drawn only once (*32*). Finally, the Maternal Brain Project includes one pregnant female participant that underwent in-vitro-fertilization (IVF) to achieve pregnancy (38 years old when onset of pregnancy). The participant delivered at full term via vaginal birth and breastfeed until 16 months postpartum. The study design included 26 sessions covering the whole pregnancy, starting 3 weeks before conception until 2 years postpartum, during which the participant underwent MRI scanning, blood draw for endocrine assessments and behavioral measurements (*29*, *30*). None of the densely sampled individuals presented any outlier for image quality, therefore all sessions were included, for a total of 30 timepoints each for the 28andMe and the 28andOC, 40 timepoints for the 28andHe and 26 timepoints for the Maternal Brain Project.

To further investigate between sexes variability, we used data from the Hangzhou Normal University (HNU) study from the Consortium for Reliability and Reproducibility (CoRR) (*35*). The HNU consists of 30 healthy adults (50% females, age mean ± SD = 24.4 ± 2.4) that underwent MRI scans 10 times over the course of one month (one scan every 3-4 days).

To further investigate brain sex in different hormonal profile, we used dense sampling data from the NeuroCycle+ study (*36*, *37*). This study included one naturally cycling 37 years old female (history of regular menstrual cycles, current use of nicotine, no history of pregnancy or breastfeeding), one 31 years old female participant taking oral contraceptives (0.03-mg ethinyl-estradiol, 2-mg dienogest, Maxim, Jenapharm) prescribed approximately 3 months before starting the study, and one male participant (36 years old) that underwent testing 25 times over 5 consecutive weeks (mostly on weekdays).

To test cycle associations in bigger sample sizes, we used data from the Hormone Health Study (HHS) (*38–40*), the HNU (*35*) and the sample described in Wright et al. (*41, 42*). The Hormone Health Study consists of 22 naturally cycling females (age mean ± SD = 21.47 ± 3.11) that underwent MRI scan during menses, ovulation and mid-luteal phase. Blood samples were collected at each measurement to assess the levels of hormones (*38–40*). We excluded 7 participants for which hormonal levels did not match the estimated phase of the cycle. The sample described in Wright et al. included 21 healthy females (age mean ± SD = 22.98 ± 3.58) tested at three timepoints across the menstrual cycle (early follicular phase - EFP, late follicular phase – LFP, and mid-luteal phase - MLP). Phases were assessed based on self-reported onset day of menses as: EFP → days 1-4 of a cycle, LFP → days 10-12 and MLP → days 20-22. Hormone samples were collected at each timepoint to confirm the estimated phase of the cycle (*41, 42*). 18 participants for which the estimated cycle phase did not correspond to hormone levels were excluded from the analysis. For the HNU we selected female participants and used the variable “last menstrual period” to assign each session to EFP, LFP and MLP based on the same criteria as Wright and colleagues.

### Image segmentation and features selection

Raw T1-weighted MRI images were preprocessed in FreeSurfer (v. 7.1.1) and automated cortical and subcortical reconstruction was carried out. We used the Euler number, a proxy of structural image quality (*33*), for quality control of images and excluded outliers (i.e. values lower than three standard deviations from the mean) for the average Euler number across hemispheres. To obtain a better definition of the limbic system, we combined different segmentation approaches, as previously described (*23*, *24*), including the multimodal parcellation of the cerebral cortex (*58*), the classical FreeSurfer subcortical segmentation, the segmentation of subcortical limbic structures (*59*) and the subfield segmentation of the hippocampus (*60*) amygdala (*61*) and thalamus (*62*). The final feature set includes 493 cortical and subcortical volumes (whole brain), carefully combined to avoid overlap between regions derived with different segmentation approaches, of which 160 features were assigned to the limbic feature set (see Supplementary Table T1 for the complete list) and 333 regions were assigned to the non-limbic feature set. Supplementary Figure S18 depicts the distribution of the cortical limbic and non-limbic volumes.

Correction for total brain size, was performed in the training sample by matching each female and male participant according to their eTIV difference, ensuring a difference in eTIV of less than 3% and a final training sample with no significant difference in eTIV between sexes.

### Sex classification models and individual’s trajectories

Binary classification models implemented in the *xgboost* (version 1.7.5.1) package in R were trained for each set of features, as previously described in Matte Bon et al. (*23*). Sex was binary coded in the training sample, with 0 assigned to males and 1 assigned to females. The training sample was balanced by sex, age, eTIV and Euler number. The classification model consisted of a nested 5-fold cross validation, with initial number of rounds set to 1000 and learning rate *η*=0.01. Cross-validation folds were stratified by sex in the outer loop of the nesting procedure. To determine the optimal number of iterations to use for training the final models on the full set of data, the prediction error of the inner loop was assessed at each iteration. The resulting class probabilities had a range between 0 (*male-like* brain) and 1 (*female-like* brain). Label threshold for establishing model performance was set at 0.5, with values below 0.5 binary labeled as males and values higher than 0.5 binary labeled as females. Model performance was assessed by computing the area under the receiving operating characteristic curves (AUC-ROC) and ranged between 0.817 (limbic model) and 0.880 (whole brain) and the accuracies ranging between 75% and 81% (limbic and whole brain models respectively) (see Supplementary Table T3 for details). Feature contribution to the models was assessed using the xgb.importance function. For each regional model the feature gains were extracted and compared to the feature gains of the whole brain model (Supplementary Figure S2). A supplementary analysis stratifying the cross-validation folds also in the inner loop yielded similar estimates, with correlation coefficients > .91 for all models (Supplementary Figure S19).

To independently validate our models in external samples, we applied the trained models to the 28andMe, 28andOC, 28andHe and Maternal Brain Project. Model validation ensured independence of sex prediction between observations by predicting at the individual observation level from a model trained in a large, independent dataset. We then stored the obtained class probabilities and looked into trajectories of change for each dataset across timepoints, particularly focusing on hormonal states. For this purpose, sessions were divided for the naturally cycling female of the 28andMe and NeuroCycle+ data according to the phase of the cycle into menses, follicular phase, ovulation window and luteal phase, while pregnancy was divided into pre-conception, first trimester, second trimester, third trimester and postpartum. Trajectories across hormonal states were then compared between subjects.

### Statistical analysis

All analyses were performed in R (version 4.3.1). To assess to which degree different training sample characteristics affected brain sex estimates, correlations between class probabilities obtained with different eTIV correction methods (matching, raw features, eTIV regression and PCP) were applied. To assess the degree of agreement between models run in the different sample, Cohen’s kappa was calculated using the psych package (version 2.4.1) and interpreted the results according to the literature defining “slight” (0.00 – 0.20), “fair” (0.21 – 0.40), “moderate” (0.41 – 0.60), “substantial” (0.61 – 0.80) and “almost perfect” (0.81 – 1.00) (*34*).

Trajectories of class probabilities were plotted for each dataset according to the hormonal state during data collection. For the naturally cycling female, sessions were divided according to the phase of the cycle into menses, follicular phase, ovulation window and luteal phase. Importantly, because the start of the experiment was blinded according to the phase of the cycle, the first experimental session was carried out in the luteal phase, resulting in data collection spanning across consecutive two menstrual cycles (luteal phase to luteal phase). Therefore, when dividing data into cycle phases, the luteal phase included data from two consecutive menstrual cycle. For the pregnant participant, time points were divided into pre-conception, first trimester, second trimester, third trimester and postpartum.

To investigate the individual’s trajectories of brain sex across hormonal states, we first performed a Levene’s test for equality of variances to investigate differences in variance distribution in class probabilities between subjects, using the *car* (version 3.1-2) package. Since the distribution of class probabilities obtained with each model are skewed toward the biological sex of the participants, we centered each distribution around 0 by subtracting the mean for each subject from each observation. To control for multiple comparisons, we applied Bonferroni correction, considering a total of 18 comparisons (six comparison for each model), establishing the new significance threshold at alpha = 0.0028. A supplementary analysis using the same approach was carried out in the NeuroCycle+ sample. Since no pregnant individual was present in this sample, we considered a total of 9 comparisons (three for each model) and established the Bonferroni correction threshold at alpha = 0.005. To exclude pre-existent univariate variance differences that could affect the variance of the predictions, we further investigated difference in variance at a univariate level, comparing the coefficient of variation obtained for limbic and non-limbic features between the naturally cycling female and the male participant. To further explore whether patterns of higher variance in one sex were present in the training sample, we applied a Levene’s test to test for differences in variance of cross-sectional class probabilities between males and females in the training sample. We applied Bonferroni correction for multiple comparison, considering three models and establishing the new threshold at alpha = 0.0167.

To test for within subject variance differences between sexes in the HNU sample, we applied for each machine learning model two linear mixed effects models (LME) using the *nlme* package (version 3.1-165), accounting for sex, session, age and Euler number, one considering equal variance between sexes and one allowing different variance by introducing a weight term. We then performed an anova to statistically compare the models. Significant results indicate different variance between sexes.

To compare estimates of brain sex across different hormonal states, we applied a pairwise t-test between naturally cycling and oral contraceptive use in the same female subject. Correction for three comparisons (one for each model) was considered using Bonferroni.

Correlations between class probabilities and hormonal levels were performed for each subject across timepoints. Hormones were selected based on data availability, excluding days with missing values for each hormone. For the naturally cycling and oral contraceptive-using female, concentrations of estradiol, progesterone, LH and FSH were collected. For the pregnant female, estradiol, progesterone and LH were assessed. Finally, the saliva concentration of total testosterone and cortisol and the serum levels of total testosterone, free testosterone, estradiol and cortisol were assessed for the male participant. Bonferroni correction for multiple comparison was carried out with the threshold established based on the number of hormones available for each participant (12 comparisons for the naturally cycling and oral contraceptive user, 9 comparisons for pregnancy and 18 comparisons for the male participant).

Finally, as a follow up analysis based on the correlation results, to test temporal dependencies between class probabilities and estradiol levels in the naturally cycling female in the limbic and whole brain model, a vector autoregressive model was applied. We used the *vars* package (version 1.6-1). Based on similar models in the same data (*16*) we first considered a second-order model with lag = 2 for consistency. We then applied the function *VARselect* in the *vars* package to estimate the best lag based on the information criteria for sequential increasing the lag order and we ran also a third-order VAR model with lag = 3. Final models had the following structure:

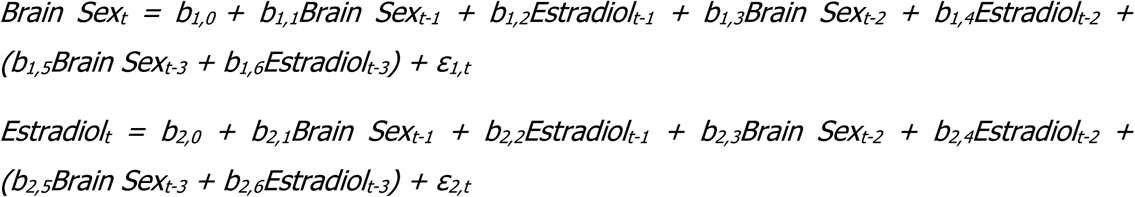

where ε represents the error terms and the terms in parentheses are added only for the third-order VAR models. When significant associations emerged, we applied a Granger causality test using the *causality* function in the same package.

## Acknowledgments

This work was performed as part of the International Research Training Group: Women’s Mental Health Across the Reproductive Years (IRTG 2804). The study was supported by the BMBF-funded de.NBI Cloud within the German Network for Bioinformatics Infrastructure (de.NBI) (031A537B, 031A533A, 031A538A, 031A533B, 031A535A, 031A537C, 031A534A, 031A532B). TK received funding from the Interfaculty Graduate Program AI4Med-BW and Fortüne Program (2660-0-0) from the Faculty of Medicine at University of Tübingen, the German Research Foundation (IRTG 2804), the Research Council of Norway (#323961) and the European Research Council (ERC CoG, #101086793, HealthyMom). TK is a member of the Machine Learning Cluster of Excellence, EXC number 2064/1 – Project number 39072764. BD received funding from the German Research Foundation (IRTG 2804, DE2319/9-1). EC received funding from SciLifeLab and the Swedish Research Council.

## Author contributions

Conceptualization: GMB, TK. Methodology: GMB, TK. Formal analysis: GMB. Data curation: GMB, TK. Data interpretation: GMB, ACSK, TK. Visualization: GMB. Writing—Original Draft: GMB, TK. Writing—Review and Editing: GMB, ACSK, EC, BD, TK. Supervision: TK, EC, BD. Funding Acquisition: TK, EC, BD.

## Competing interests

The authors declare no competing interests.

## Data and materials availability

All data used to reach the conclusions presented in this paper are presented in the main text or the supplementary material. All raw data is available via dedicated data use agreements with the data providers (see below).

We used data from the Human Connectome Project - Young Adult (HCP-YA), the HCP-Aging (HCP-A), the Queensland Twin IMaging (QTIM), the 28andMe, the Maternal Brain Project, the 28andHe, the Hangzhou Normal University study (HNU) from the Consortium for Reliability and Reproducibility (CoRR) initiative, the NeuroCycle+, the Hormone Health Study (HHS) and the *Endocrine modulation of cortical and retinal baseline blood flow across the menstrual cycle* study (Wright et al., 2025). For the HCP-YA and HCP-A, data were provided [in part] by the Human Connectome Project, WU-Minn Consortium (Principal Investigators: David Van Essen and Kamil Ugurbil; 1U54MH091657) funded by the 16 NIH Institutes and Centers that support the NIH Blueprint for Neuroscience Research; and by the McDonnell Center for Systems Neuroscience at Washington University. QTIM data used in the preparation of this article were obtained from the Queensland Twin IMaging (QTIM) study. The QTIM study was supported by the National Institute of Child Health and Human Development (R01 HD050735), and the National Health and Medical Research Council (NHMRC 496682, 1009064), Australia. Access to CoRR data was obtained via the International Neuroimaging Data-sharing Initiative (INDI). The National Institute on Drug Abuse (NIDA) and the National Natural Science Foundation of China (NSFC) have been instrumental in the CoRR collaboration providing the necessary funding and manpower to build the foundation of the project along with the Child Mind Institute, the Institute of Psychology, Chinese Academy of Sciences and the Nathan Kline Institute. Access to QTIM, 28andMe, 28andHe, the Maternal Brain Project, HHS and NeuroCycle+ was available throughout OpenNeuro. Data from Wright et al., 2025 were accessible throughout G-node Open Data.

## Code availability

All code will be made publicly available at https://github.com/gloriamatte upon publication.

## Notes

### Competing Interest Statement

The authors have declared no competing interest.

### Summary of Updates

Addition of new data and supplementary analysis, revisions of Figures, addition of several new supplementary figures and tables

