## Supplementary Figures S1-S19 and Supplementary Table T1-T3 for "Inter- and intra-individual trajectories of brain sex in different neuroendocrine profiles"

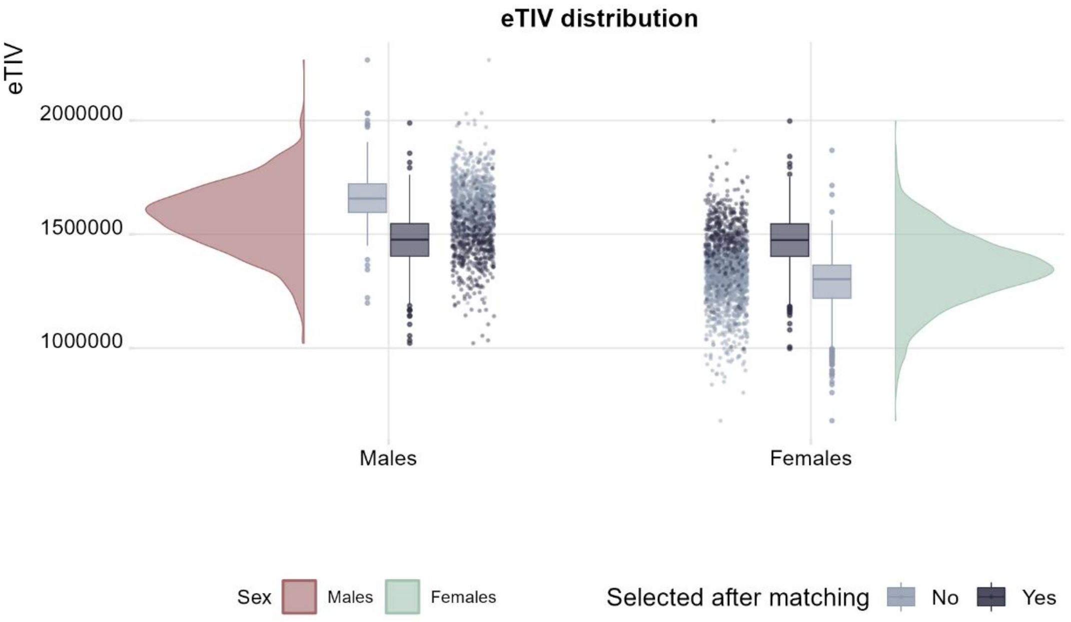


**Figure S1. Estimated total intracranial volume (eTIV) distribution of the matched sample compared to the general population.** The eTIV distribution of the selected participants largely overlapped with the general sample distribution for each sex, ensuring that the final selected sample after matching included some individuals toward both ends of the distribution.

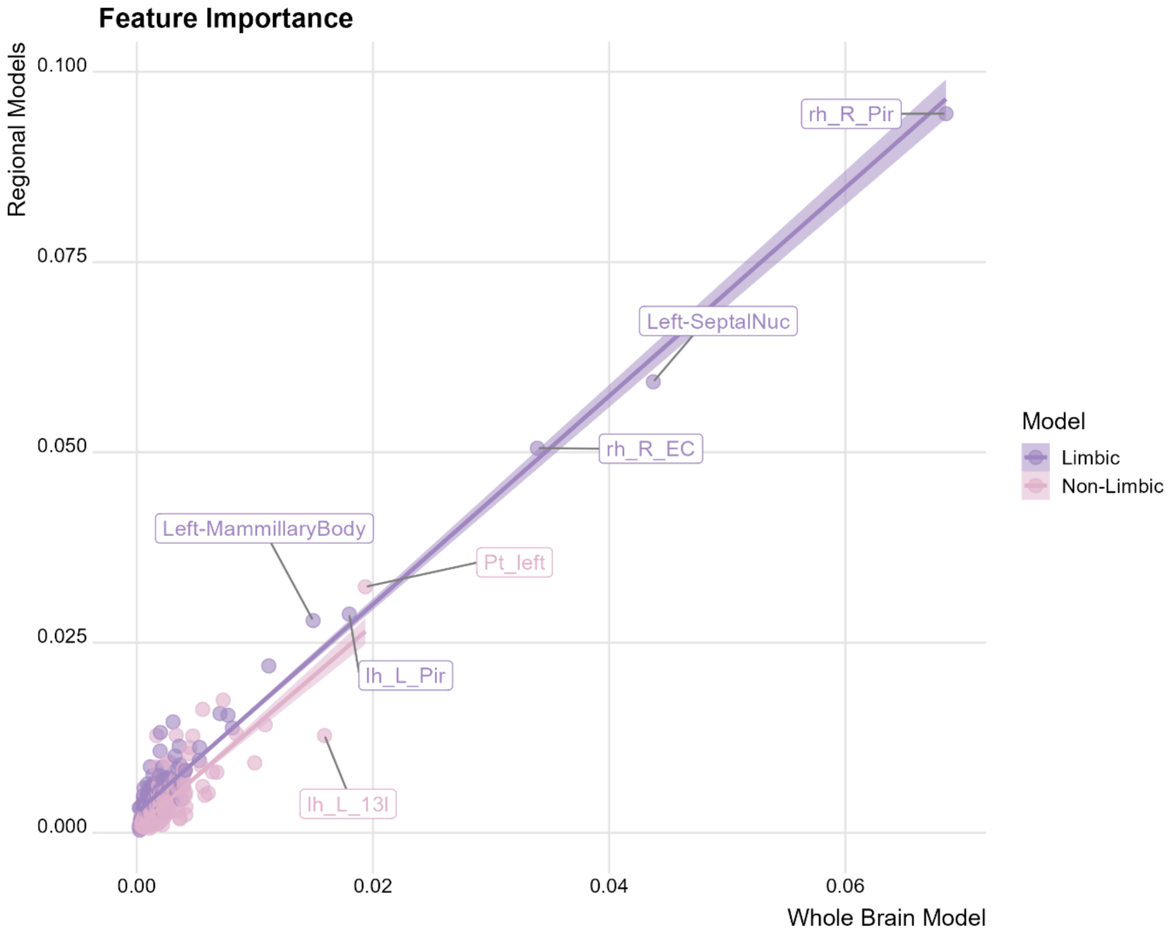


**Supplementary Figure S2. Correlation between feature gains for the regional and whole brain models.** Features contributing the most to the two regional models also contributed the most to the whole brain model. (Abbreviations: EC: Entorhinal cortex, Pir: Piriform Cortex, SeptalNuc: Septal Nuclei, Pt: Paratenial thalamic nuclei).

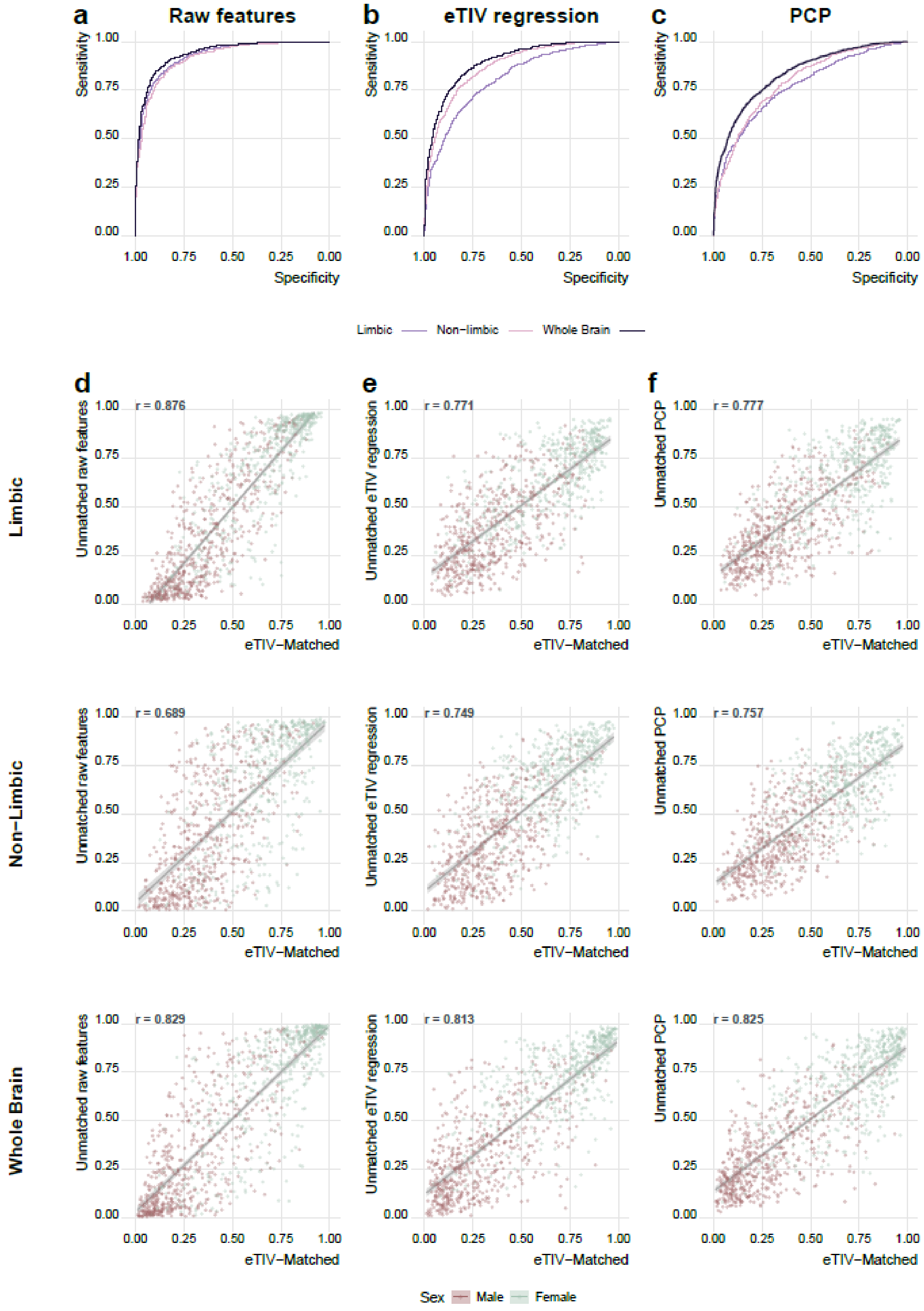


**Supplementary Figure S3. Models trained on a bigger sex-balanced sample yield similar predictions to the models trained on the eTIV-matched sample. a)** Models trained on raw features without eTIV correction methods reached higher performances. **b-c)** Models corrected for eTIV with regression or power-corrected proportion (PCP) method yielded performances similar to the models trained on our eTIV-matched sample despite the increase in sample size. **d-f)** Class probabilities between the eTIV-matched sample and the sex-balanced sample were significantly correlated for all models.

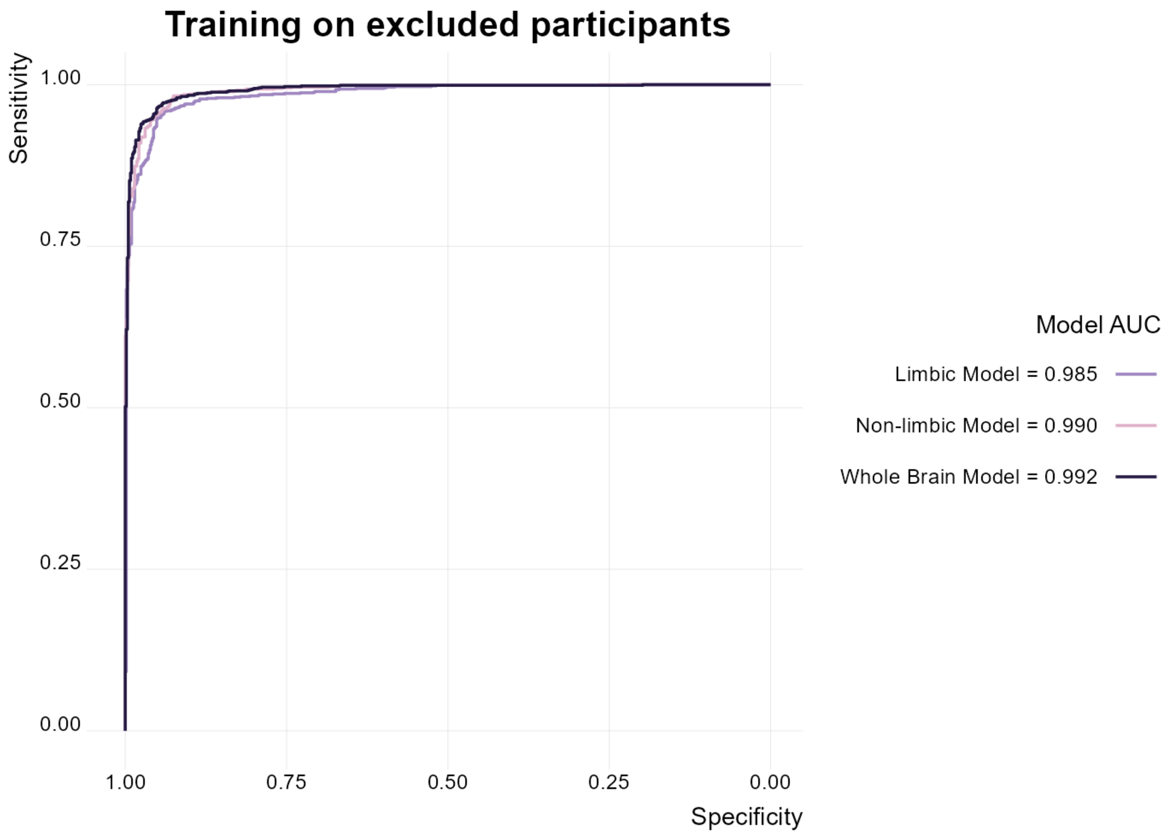


**Supplementary Figure S4. Models trained on the participants excluded by the eTIV-matching procedure yielded highly biased prediction.** All models achieved almost perfect classification, with AUCs of over .98.

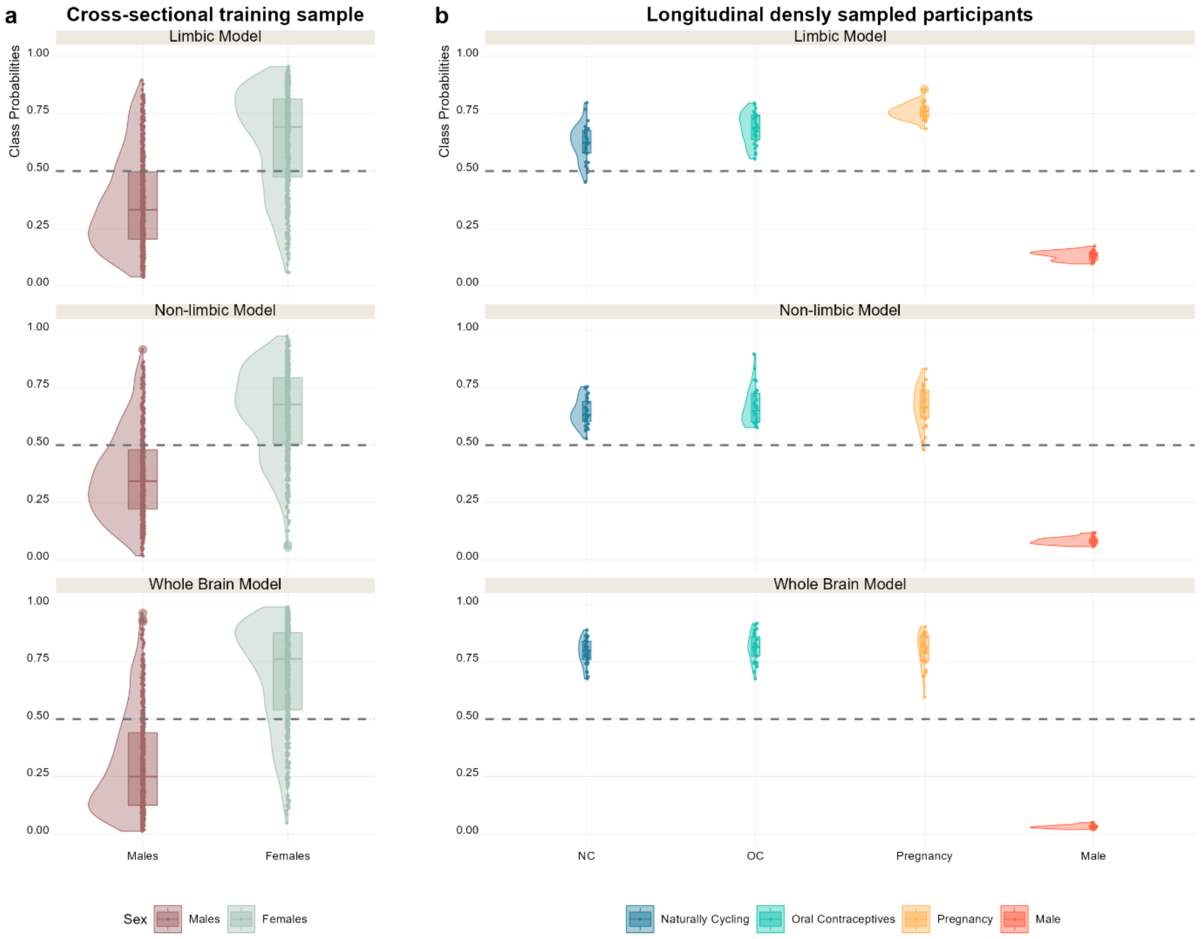


**Supplementary Figure S5. Class probabilities distribution in the training and independent test sets. a)** Class probabilities for male and female participants obtained by cross-validation in the training sample. **b)** Distribution of class probabilities in the longitudinal densely sampled individual, showing correct classification labels according to their biological sex for all participants for most timepoints. Data points below the dotted line are binary labeled by the model as "male", while data points above the dotted line are labeled as "female".

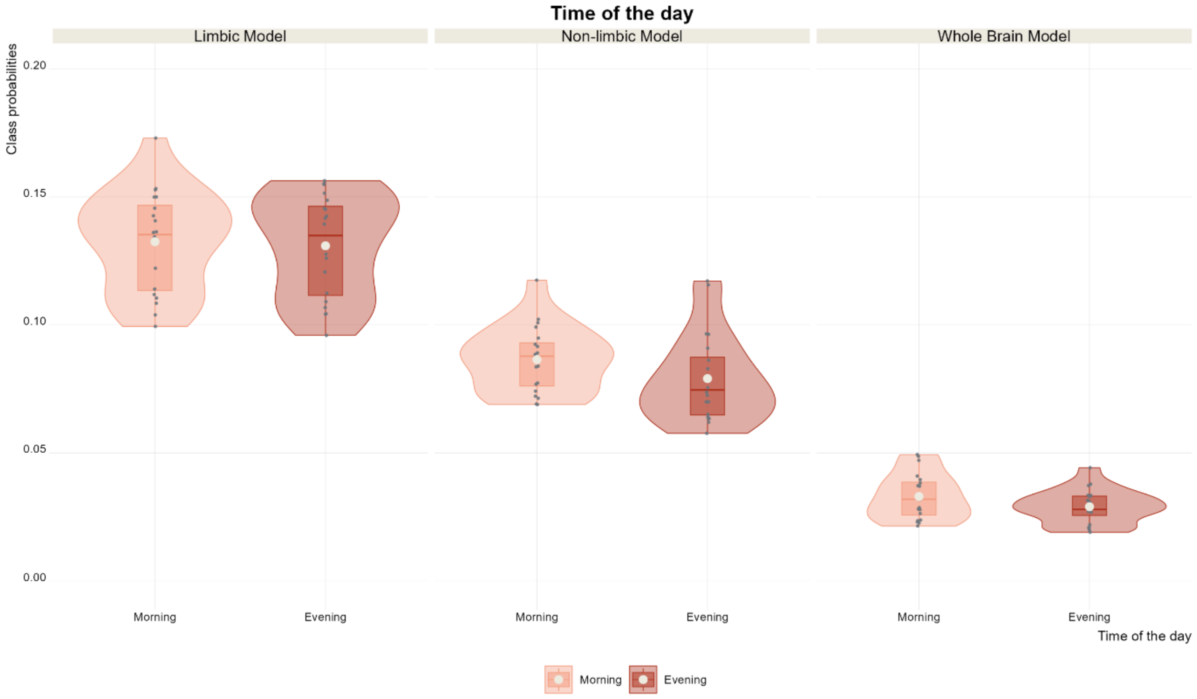


**Supplementary Figure S6. Distribution of class probabilities in the male participant by time of the day.** No significant difference between morning and evening sessions was found for any model.

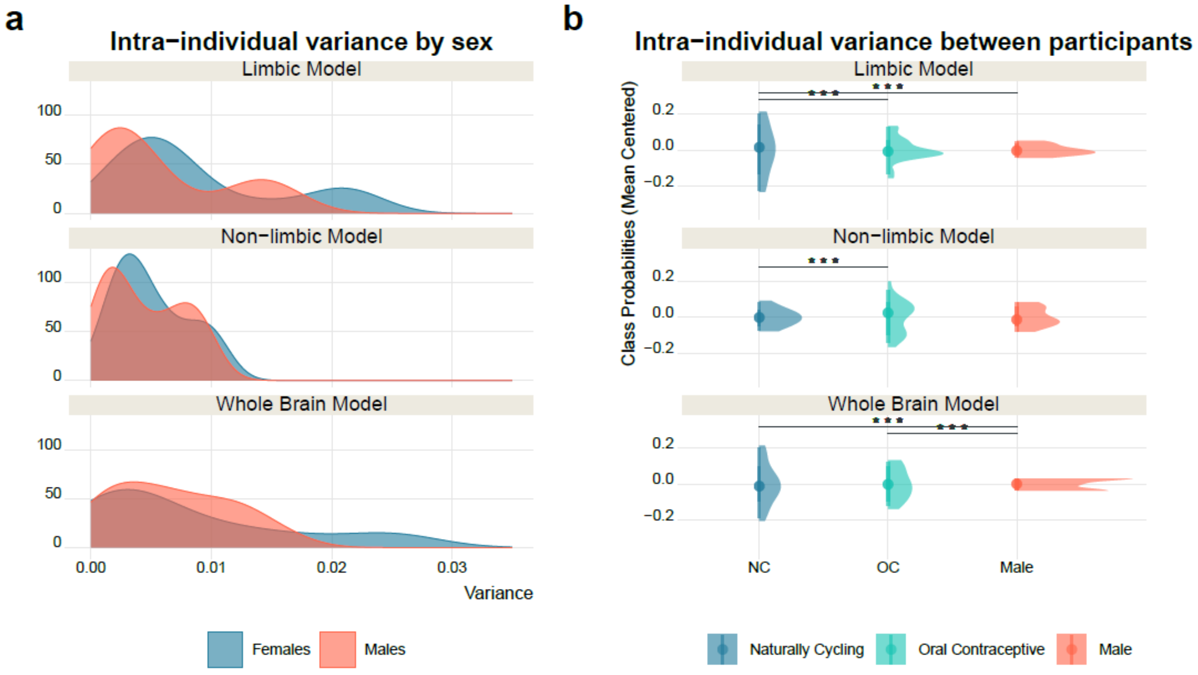


**Supplementary Figure S7. Variance distributions of class probabilities between sexes. a)** Pattern of higher variability over time in brain sex estimates in females compared to males in data from the HNU study. **b)** Independent densely sampled participants from the NeuroCycle+ study show the same pattern of higher variance in the naturally cycling female compared to the male in the limbic and whole brain model. *** p < .005 (Bonferroni-corrected threshold)

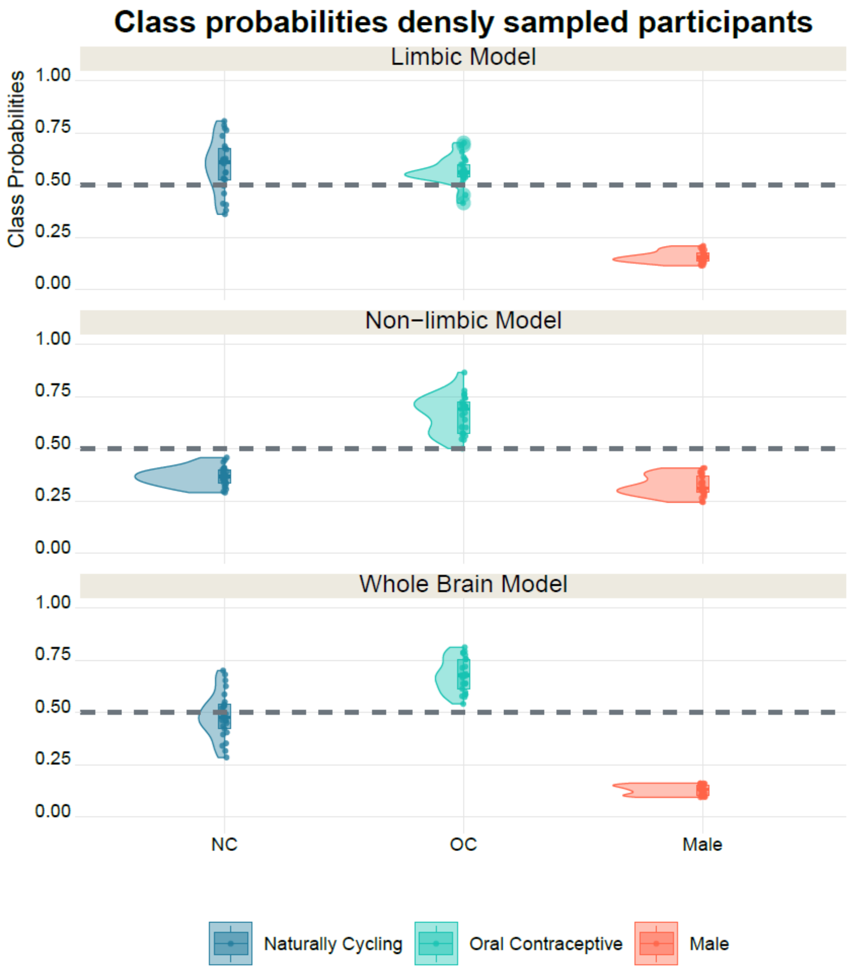


**Supplementary Figure S8. Distribution of class probabilities of densely sampled participants in the NeuroCycle+ study.** The non-limbic model classified the naturally cycling female participant as male-like for all timepoint, potentially masking any difference in variance with the male participant. Dotted line represents the 0.5 threshold for binary classification into male-like (<0.5) and female-like (> 0.5).

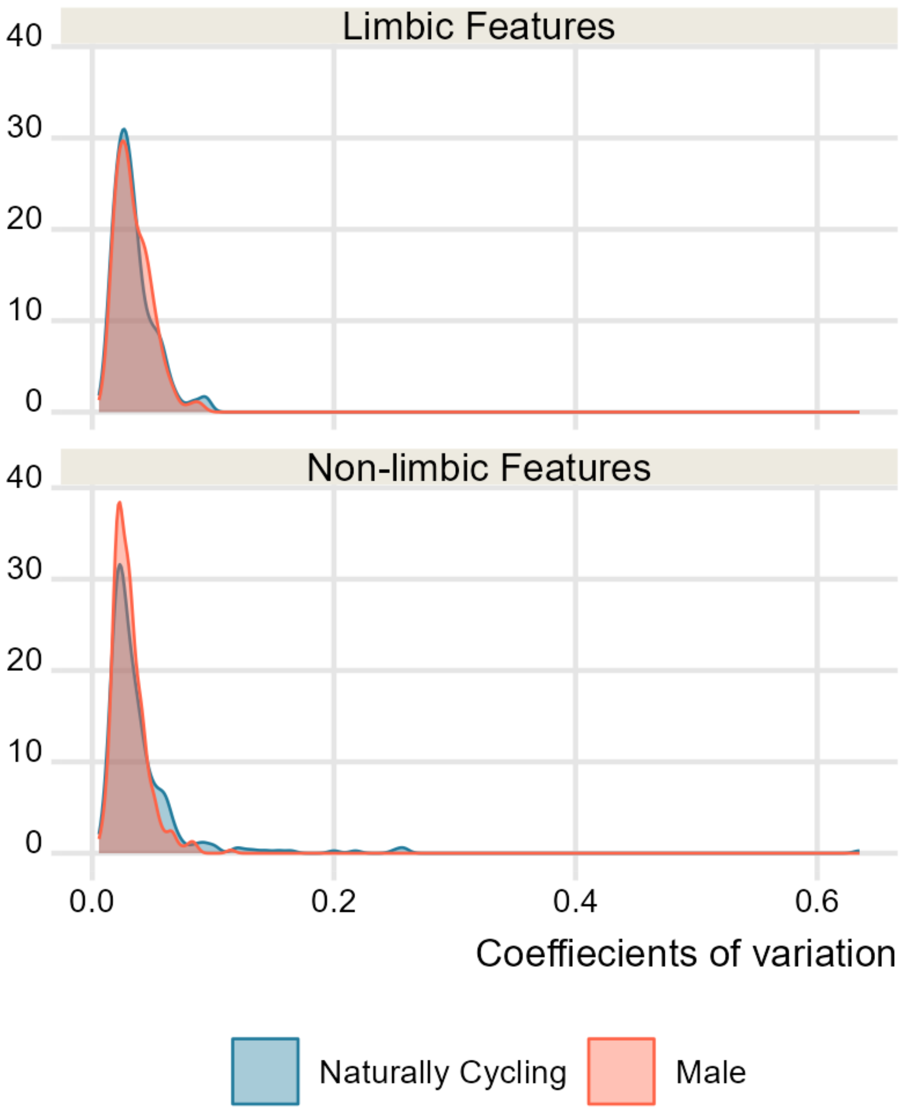


**Supplementary Figure S9. Distribution of the univariate coefficients of variation overlapped between the naturally cycling female and the male.** Coefficients of variation were calculated as a measure of variability for each region across time points in each subject. Density plots represent the distribution of all limbic and non-limbic coefficients of variation. Limbic and non-limbic volumes showed overall similar variability between individuals, suggesting that the lower variance in class probabilities showed by the male subject is not attributable to lower variance in anatomical data.

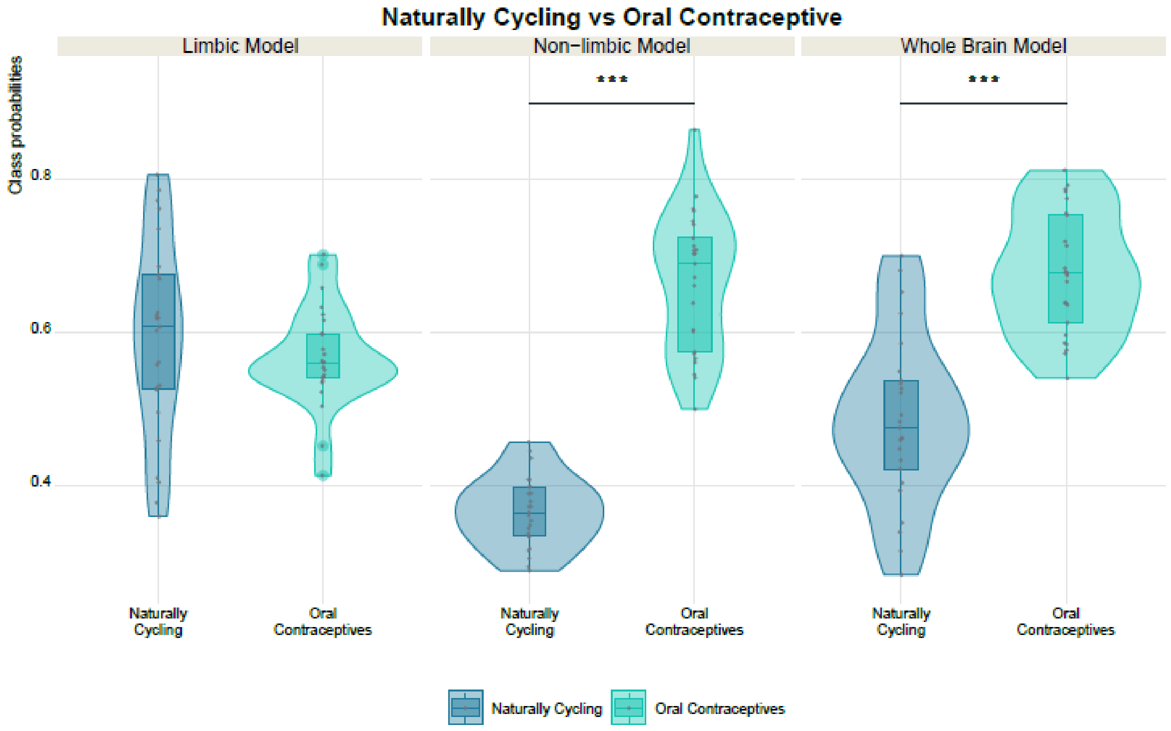


**Supplementary Figure S10. Patterns of more female-like brain in the OC user compared to the naturally cycling participant in the NeuroCycle+ sample.** Significantly higher class probabilities were found for the non-limbic and whole brain model. *** p < .017 (Bonferroni-corrected threshold)

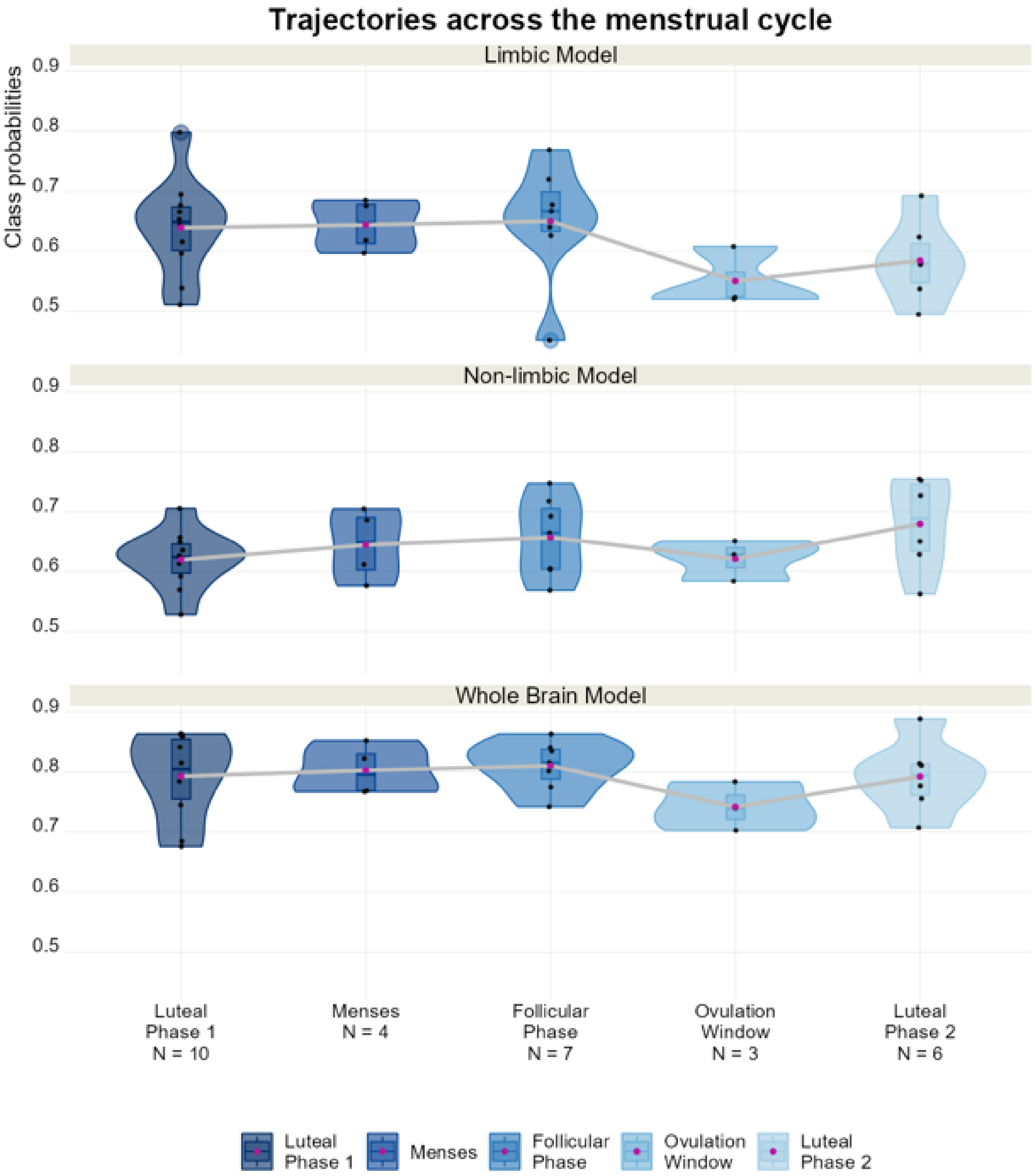


**Supplementary Figure S11. Trajectories of brain sex across the two consecutive menstrual cycles showed negative peak around ovulation.** Data collection started during the luteal phase (cycle 1) and lasted until the mid-luteal phase of the following one (cycle 2), partially covering two consecutive menstrual cycles.

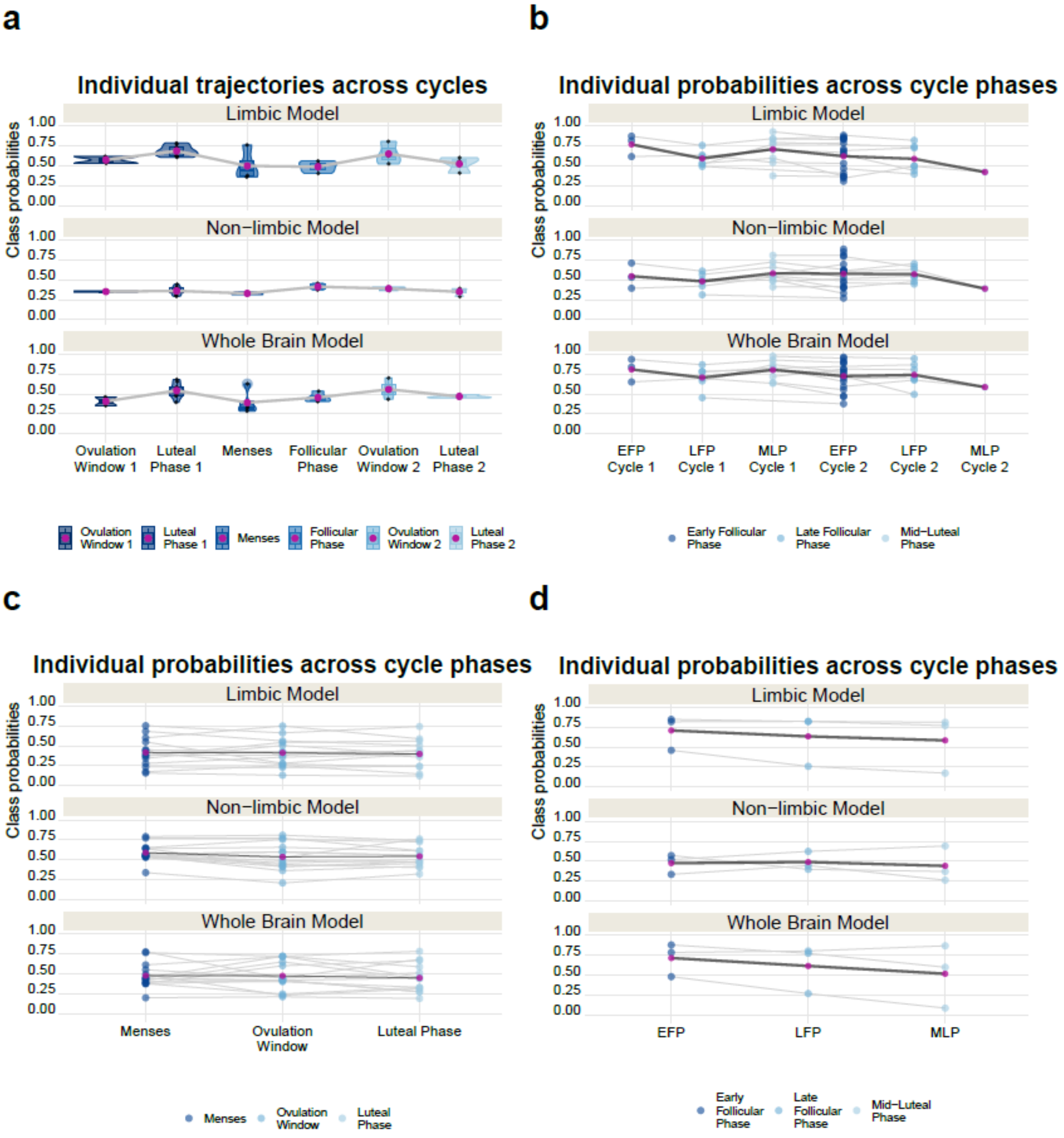


**Supplementary Figure S12. Inter- and intra-individual dynamics of brain sex across menstrual cycle phases.** Consistent trajectories between models across menstrual cycle phase were found in the naturally cycling participant from the NeuroCycle+ sample, with a pattern of increase brain sex estimates in the second half of the cycle (luteal phase 1, ovulation window 2) in two consecutive cycles **(a).** Pattern of inter- and intra-individual variability across menstrual cycle phases emerged in the HNU **(b)**, HHS **(c)** and Wright et al. samples **(d)** for all models.

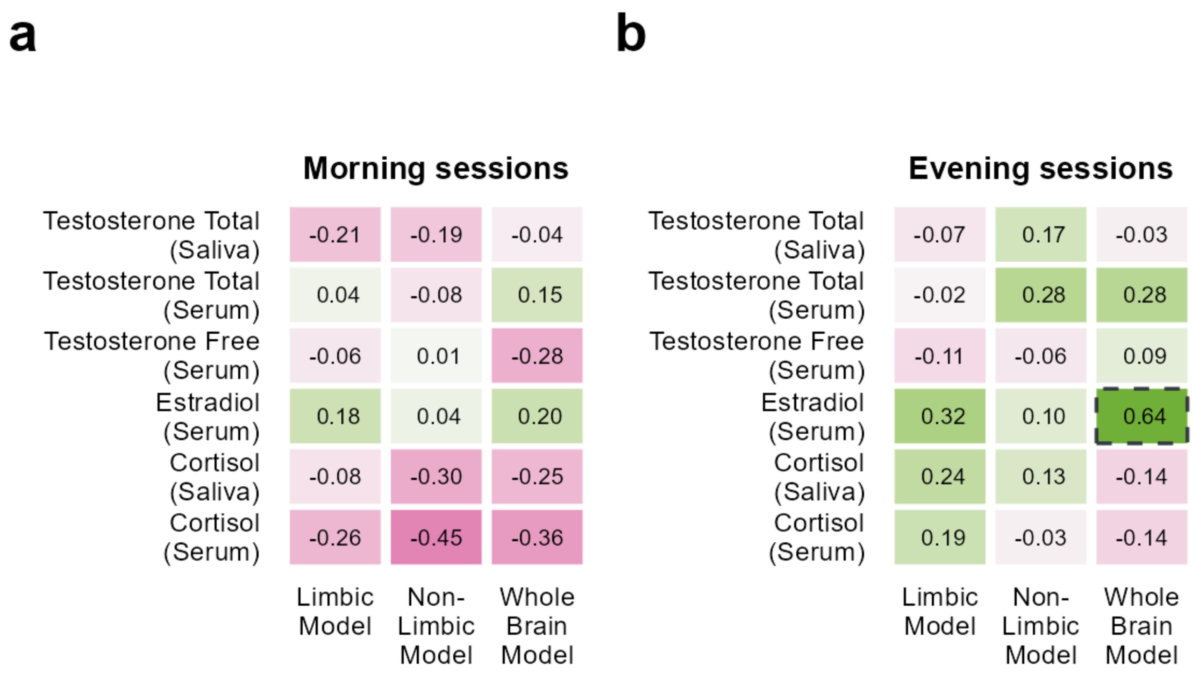


**Supplementary Figure S13. Evening estradiol levels are positively correlated with whole brain estimates in the male individual. a)** Correlations of class probabilities with hormone levels across the morning sessions in the male participant showed no significant associations. **b)** Across evening sessions, a nominally significant positive correlation between whole brain class probabilities and estradiol levels was found that did not survive Bonferroni correction for multiple comparisons. Pink color indicates negative correlation, while green color indicates positive correlation. Dashed lines: $p < .05$

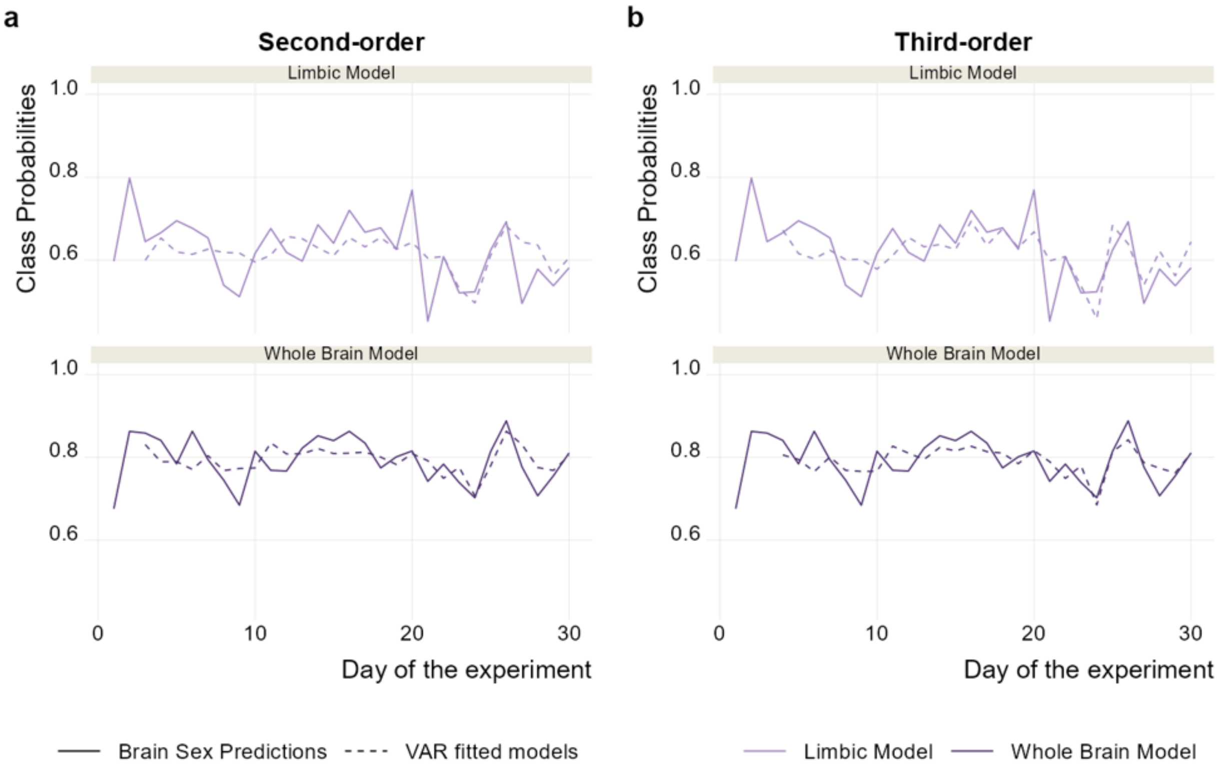


**Supplementary Figure S14. Limbic class probabilities are associated with previous estradiol levels. a)** Comparisons between brain sex prediction and fitted values for second-order VAR models. b) Comparisons between brain sex predictions and fitted values for third-order VAR models. Menses: experimental days 11 to 14; ovulation: experimental days 22 to 24.

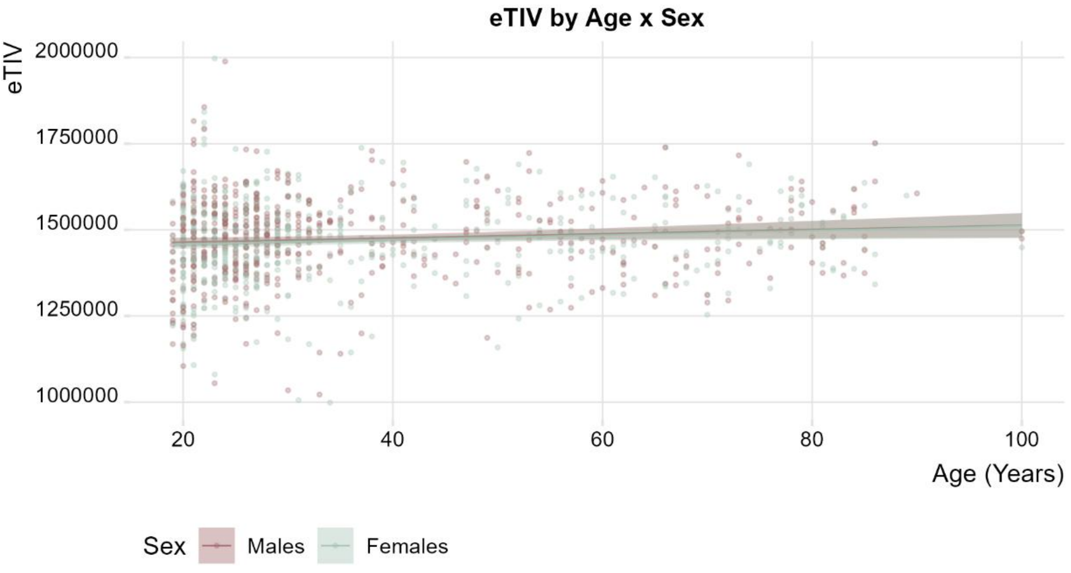


**Supplementary Figure S15. eTIV of the matched sample stratified by age and sex.** When considering potential bias due to an interaction effect of age and sex on eTIV, our matched training sample showed no significant interaction effect ($t = 0.128$, $p = .898$), ensuring that physiological changes in eTIV due to the aging process, such as volume loss, will not bias the classifier toward one of the two sexes.

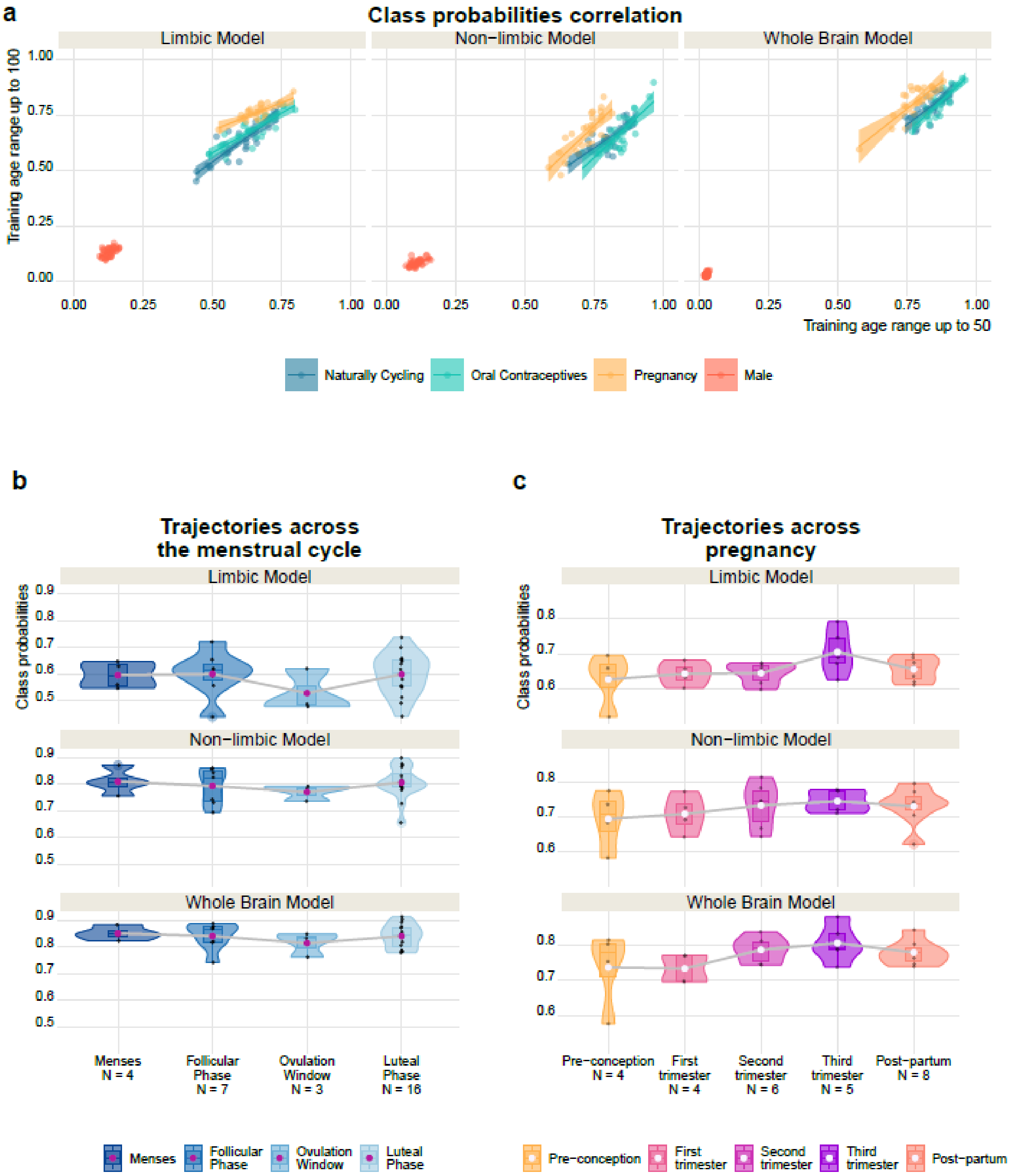


**Supplementary Figure S16. Model training in a closer age range to densely sampled participant yielded similar brain sex dynamics for all participants. a)** Class probabilities obtained training in an eTIV-matched sample with maximum age of 49 or 100 years old are correlated for all participants. **b)** Trajectories of brain sex across menstrual cycle phases show a dip around ovulation. **c)** Trajectories across pregnancy show a maximum in the third trimester.

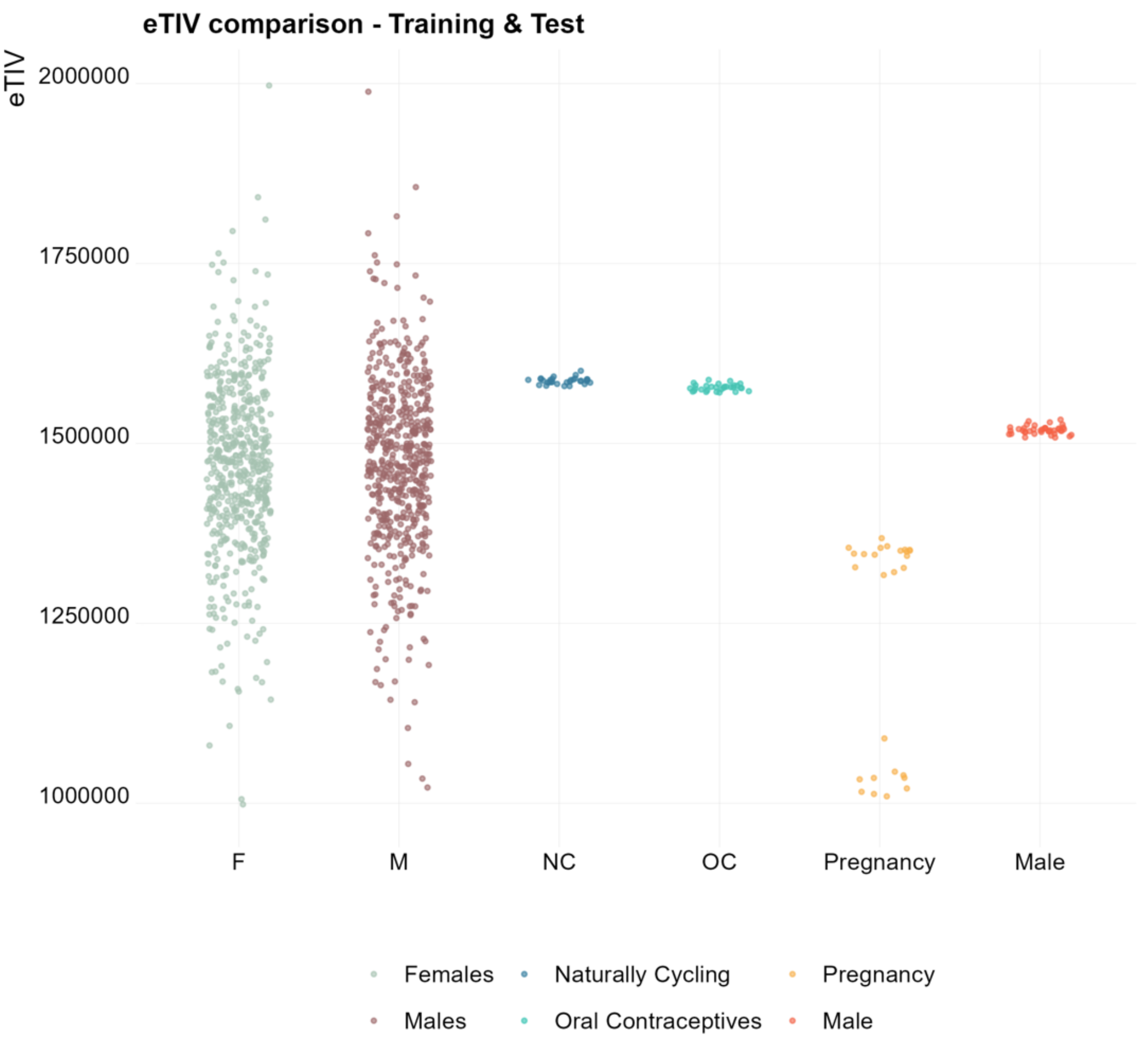


**Supplementary Figure S17. eTIV distribution in the densely sampled individuals compared to the eTIV-matched training sample.** The eTIV of all participants overlap with the distribution in the training sample, assuring that the matching procedure was representative also of the independent test sets.

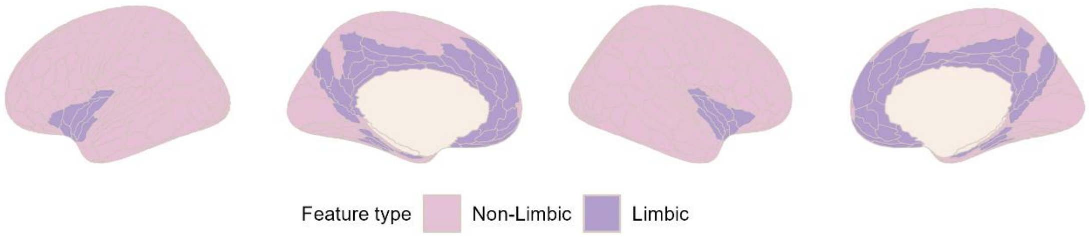


**Supplementary Figure S18. Visualization of the cortical limbic and non-limbic regions.** A complete list of cortical and subcortical structures included in the limbic model is available in the Supplementary Table T1.

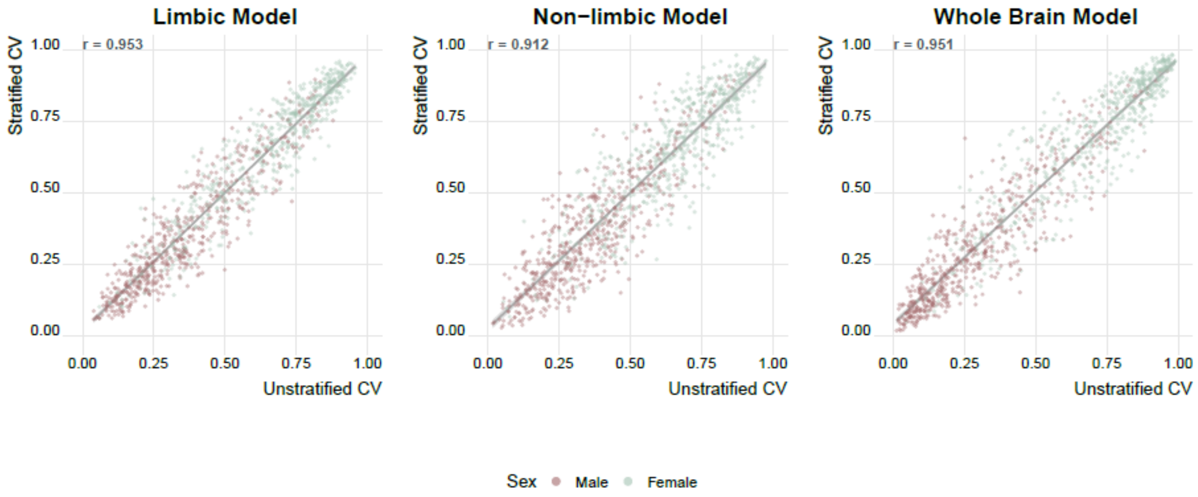


**Supplementary Figure S19. Correlation of class probabilities obtained with sex-stratification in the outer or inner and outer loop of the nested cross-fold validation.** Both training procedures yielded similar estimates, with correlation coefficients > .90 for all models.

**Supplementary Table T1. List of limbic structures with the correspondent feature in Freesurfer**

| Structure | FreeSurfer Feature | Segmentation |
|---|---|---|
| Anterior Cingulate Cortex | 33pr | Glasser atlas |
| | p24pr | |
| | a24pr | |
| | p24 | |
| | a24 | |
| | p32pr | |
| | a32pr | |
| | d32 | |
| | p32 | |
| | s32 | |
| | 8BM | |
| | 9m | |
| | 10v | |
| | 10r | |
| | 25 | |
| Orbitofrontal Cortex | OFC | |
| | pOFC | |
| Insula | MI | |
| | AVI | |
| | AAIC | |
| | Ig | |
| | PI | |
| | PoI1 | |
| | PoI2 | |
| Piriform Cortex | Pir | |
| Entorhinal Cortex | EC | |

| Structure | FreeSurfer Feature | Segmentation |
|---|---|---|
| Parahippocampal Area | PHA1 | Hippocampal Subfields |
| | PHA2 | |
| | PHA3 | |
| Posterior Cingulate Cortex | DVT | |
| | ProS | |
| | POS1 | |
| | POS2 | |
| | RSC | |
| | v23ab | |
| | d23ab | |
| | 31pv | |
| | 31pd | |
| | 31a | |
| | 23d | |
| | 23c | |
| | PCV | |
| Hippocampus | Hippocampal_tail | |
| | subiculum-body | |
| | CA1-body | |
| | subiculum-head | |
| | hippocampal-fissure | |
| | presubiculum-head | |
| | CA1-head | |
| | presubiculum-body | |
| | parasubiculum | |
| | molecular_layer_HP-head | |
| | molecular_layer_HP-body | |
| | GC-ML-DG-head | |

| Structure | FreeSurfer Feature | Segmentation |
|---|---|---|
| | CA3-body | |
| | GC-ML-DG-body | |
| | CA4-head | |
| | CA4-body | |
| | fimbria | |
| | CA3-head | |
| | HATA | |
| Amygdala | Lateral-nucleus | Nuclei of Amygdala |
| | Basal-nucleus | |
| | Accessory-Basal-nucleus | |
| | Anterior-amygdaloid-area-AAA | |
| | Central-nucleus | |
| | Medial-nucleus | |
| | Cortical-nucleus | |
| | Corticoamygdaloid-transitio | |
| | Paralaminar-nucleus | |
| Anterior and Dorsomedial Thalamic Nuclei | AV | Thalamic Nuclei |
| | LD | |
| | MDl | |
| | MDm | |
| Nucleus Accumbens | Nucleus-Accumbens | ScLimbic |
| Hypothalamus | HypoThal-noMB | |
| Fornix | Fornix | |
| Mammillary Body | MammillaryBody | |
| Forebrain | Basal-Forebrain | |
| Septal Nuclei | SeptalNuc | |

**Supplementary Table T2. Cohen's Kappa values indicate the agreement between eTIV matched models and models trained on the sex-stratified sample with different eTIV correction methods.**

**Limbic Model**

| | eTIV regression | PCP | Raw features |
|---|---|---|---|
| **eTIV-Matched** | 0.61 | 0.62 | 0.72 |
| **eTIV regression** | 1.00 | 0.77 | 0.54 |
| **PCP** | | 1.00 | 0.56 |
| **Raw features** | | | 1.00 |

**Non-Limbic Model**

| | | | |
|---|---|---|---|
| **eTIV-Matched** | 0.56 | 0.58 | 0.49 |
| **eTIV regression** | 1.00 | 0.62 | 0.48 |
| **PCP** | | 1.00 | 0.44 |
| **Raw features** | | | 1.00 |

**Whole Brain Model**

| | | | |
|---|---|---|---|
| **eTIV-Matched** | 0.68 | 0.67 | 0.66 |
| **eTIV regression** | 1.00 | 0.74 | 0.56 |
| **PCP** | | 1.00 | 0.54 |
| **Raw features** | | | 1.00 |

**Supplementary Table T3. Model performances in the training set.** The table depicts the accuracy, sensitivity and specificity of the three models.

| | Accuracy (95% CI) | Sensitivity | Specificity |
|---|---|---|---|
| **Limbic Model** | 0.7330 (0.7057, 0.7591) | 0.7523 | 0.7138 |
| **Non-Limbic Model** | 0.7661 (0.7398, 0.7909) | 0.7706 | 0.7615 |
| **Whole Brain Model** | 0.7982 (0.7731, 0.8216) | 0.8092 | 0.7872 |
